# CaaX-motif adjacent residues control G protein prenylation under suboptimal conditions

**DOI:** 10.1101/2023.07.04.547731

**Authors:** Mithila Tennakoon, Waruna Thotamune, John L. Payton, Ajith Karunarathne

## Abstract

Prenylation is a universal and irreversible post-translational modification that supports membrane interactions of proteins involved in various cellular processes, including migration, proliferation, and survival. Thus, dysregulation of prenylation contributes to multiple disorders, including cancers, vascular diseases, and neurodegenerative diseases. During prenylation, prenyltransferase enzymes tether metabolically produced isoprenoid lipids to proteins via a thioether linkage. Pharmacological inhibition of the lipid synthesis pathway by statins has long been a therapeutic approach to control hyperlipidemia. Building on our previous finding that statins inhibit membrane association of G protein γ (Gγ) in a subtype-dependent manner, we investigated the molecular reasoning for this differential. We examined the prenylation efficacy of carboxy terminus (Ct) mutated Gγ in cells exposed to Fluvastatin and prenyl transferase inhibitors and monitored the subcellular localization of fluorescently tagged Gγ subunits and their mutants using live-cell confocal imaging. Reversible optogenetic unmasking-masking of Ct residues was used to probe their contribution to the prenylation process and membrane interactions of the prenylated proteins. Our findings suggest that specific Ct residues regulate membrane interactions of the Gγ polypeptide statin sensitivity, and prenylation efficacy. Our results also show that a few hydrophobic and charged residues at the Ct are crucial determinants of a protein’s prenylation ability, especially under suboptimal conditions. Given the cell and tissue-specific expression of different Gγ subtypes, our findings explain how and why statins differentially perturb heterotrimeric G protein signaling in specific cells and tissues. Our results may provide molecular reasoning for repurposing statins as Ras oncogene inhibitors and the failure of using prenyltransferase inhibitors in cancer treatment.

## 1. Introduction

Post-translational lipid modifications, including N-myristoylation, palmitoylation, prenylation, glycosylphosphatidylinositol (GPI) anchor addition, and cholesterol attachment, expand the functional and structural diversity of the eukaryotic proteome (1, 2). The chemical and physical properties, activities, and cellular distribution of proteins are significantly modified by the covalent attachment of a non-peptidic hydrophobic moiety to a protein. This results in the interaction of post-translationally modified proteins with cellular membranes and facilitates multiple cellular signaling pathways (2). Protein prenylation has been studied extensively due to its significance in the proper cellular activity of numerous proteins. Prenylation includes both farnesylation and geranylgeranylation and is an irreversible covalent post-translational modification found in all eukaryotic cells. These modification reactions are catalyzed by three prenyltransferase enzymes. Farnesyltransferase (FTase) or geranylgeranyltransferase type 1 (GGTase-I) catalyzes the covalent attachment of a single farnesyl (15 carbon) or geranylgeranyl (20 carbon) isoprenoid group, respectively, to a cysteine residue located in a C-terminal (Ct) consensus sequence commonly known as the “CaaX box”, in which “C” is cysteine, “a” generally represents an aliphatic amino acid, and the “X” residue determines which isoprenoid is attached to the protein target (3). Geranylgeranyltransferase type 2 (GGTase-II or Rab geranylgeranyltransferase) catalyzes the addition of two geranylgeranyl groups to two cysteine residues in sequences such as CXC or CCXX close to the Ct of Rab proteins (4, 5).

Inhibition of hepatic cholesterol biosynthesis is widely accepted to mitigate cardiovascular diseases such as Coronary Heart Disease (CHD) (6). This is achieved by inhibiting the rate-limiting enzyme, HMG-CoA reductase, of the cholesterol biosynthesis (mevalonate) pathway (7). Commonly called statins, these HMG-CoA reductase inhibitors interfere with the synthesis of intermediate products of the cholesterol biosynthesis pathway, such as isoprenoids (8, 9). Isoprenoids are precursor lipids for synthesizing cholesterol and other lipid derivatives, including farnesyl, geranyl, and geranylgeranyl pyrophosphates, squalene, dolichol, and ubiquinone. The above mentioned pyrophosphates are required for the prenylation of small and heterotrimeric G proteins (10–12). Our previous work revealed the influence of statin usage on Gβγ and, consequently heterotrimeric G-protein signaling due to the inhibition of Gγ prenylation (13). We showed that not only do statins disrupt Gγ prenylation, Gγ farnesylation is more susceptible to this inhibition than geranylgeranylation (13).

The essential role of farnesylation in modulating the oncogenic activity of Ras function led to the discovery of farnesyltransferase inhibitors (FTIs) such as Tipifarnib and Lonafarnib (14–16). The combined anti-tumor activity and low toxicity of these FTIs observed in animal models directed clinical trials to use FTIs as anti-tumor drugs (17). However, discouraging results have been observed due to the alternative prenylation exhibited in KRas and NRas (to a lesser extent), where the effect of FTIs is evaded by the activity of GGTase-I (17). Therefore, a thorough understanding of the molecular mechanism of prenylation and post-prenylation processing is crucial to develop more efficient drugs for tumor progression prevention. Given the antiproliferative effects of statins, repurposing them as anticancer drugs can also be considered (12). Observations from *in vitro* and *in vivo* pre-clinical studies and clinical studies report the anticancer effects of statins (18). Statins have shown antiproliferative effects in various cancers by primarily inhibiting the synthesis of cholesterol and its metabolites (19–21), which result in tumor growth suppression, induction of apoptosis and autophagy, inhibition of cell migration and invasion, and inhibition of angiogenesis (22–24). Therefore, it is believed that statins can influence patient survival and cancer recurrence (25). Several pleiotropic effects, including antioxidant, anti-inflammatory, and immune-modulatory properties, are also associated with statin usage, which can be causing their antiproliferative properties (26). *In vitro* studies conducted in a broad range of cancer cell lines present evidence for the anticancer properties of statins (18). For instance, Simvastatin exhibited anticancer potential in several cancer types, such as hepatoma, breast cancer, endometrial cancer, osteosarcoma, and lung adenocarcinoma (22, 27–30). Additionally, Atorvastatin exhibited anticancer effects on ovarian cancer in a study conducted using Hey and SKOV3 cells (23). Anticancer efficacy of statins has also been demonstrated during *in vivo* pre-clinical studies using xenograft animal models. For example, a xenograft mouse study showed the anti-tumor effect of Pitavastatin on glioblastoma (31). This evidence suggests the potential use of statins and prenyltransferase inhibitors as anticancer drugs in a combinatorial therapy approach. Gβγ interacts with many effectors, regulates a wide range of physiological functions, and thus has been established as a major signaling regulator (32). Although prenylation is crucial for not only Gβγ function but also GPCR-G protein signaling, molecular details are lacking on how statins influence Gγ prenylation.

Our data suggest that the peptide adjacent to the prenyl-*Cys* (pre-CaaX) region differentially regulates G protein γ sensitivity to prenylation inhibitors. Further, the described behaviors of G protein polypeptides and prenyl pyrophosphates will show how pharmacological agents, including statins and prenyltransferase inhibitors influence these proteins. Finally, our findings will further help utilize these molecular interactions to open new therapeutic windows to target G proteins-associated diseases.

## 3. RESULTS

### 3.1 Gγ subtypes show differential sensitivities to statin-mediated perturbation of the plasma membrane localization

Prenylated Gγs primarily stay bound to the plasma membrane when they are in the Gαβγ heterotrimer (Fig. 1A-Control) (33). Gγs also show a minor presence at endomembranes, likely due to heterotrimer shuttling (34, 35), activities of Guanine nucleotide Exchange Factors (GEFs) that activate minor amounts of heterotrimers (36), or interactions with endomembrane-residing GPCRs (37). Throughout this manuscript, we examined Gγ or Gγ mutant localization in HeLa cells by transfecting only fluorescently tagged Gγ subunits and relied on endogenous Gα and Gβ subunits to govern the subcellular distributions. When Gγs are not prenylated, they show a cytosolic distribution due to the failure of the membrane anchoring (13, 38). We have previously shown that while statin treatment significantly reduces the membrane anchoring of several Gγ types, indicated by their presence in the cytosol (Fig. 1-Gγ1 in control vs. Flu), other Gγ types showed a disruption of membrane binding only partially. We observed that the partial inhibition is indicated by the absence of Gγ localization at endomembranes and reduced plasma membrane localization. Further, these Gγs show minor to significant cytosolic distribution, and the extent depended on the Gγ subtype (ex: Fig. 1-Gγ2 in control vs. Flu) (13). Gγs with more cytosolic presence upon statin exposure also exhibit a nuclear localization as indicated by homogenous cell interior fluorescence. However, the molecular underpinnings of these Gγ subtype-dependent differential de-localizations of Gγ upon satin exposure were unclear. Though it is generally accepted that the CaaX motif determines the type of prenylation (farnesylation or geranylgeranylation) of a G protein, it has also been suggested that amino acids beyond this motif, extending up to 25 Ct residues, are also involved (39). To understand whether the observed statin-induced protein localization signatures of different Gγ types are exclusively dependent on their type of prenylation, we examined Fluvastatin (20 μM)-induced prenylation inhibition of the entire Gγ family (Fig. 1A-Flu). In our previous study, we tested a few statins, i.e., Fluvastatin, Lovastatin, and Atorvastatin, for their ability to disrupt membrane localization of Gγ and, thereby, Gβγ-mediated downstream signaling. Of these three statins, Fluvastatin exhibited the highest efficiency in perturbing Gγ localization and Gβγ signaling (13). Considering all the optimized conditions, we used Fluvastatin to examine statin-induced Gγ de-localization in this study. Three Gγ types reported to be exclusively farnesylated (Gγ1, 9, and 11) showed near-complete sensitivity to Fluvastatin, indicated by the cytosolic and nuclear distribution of YFP-Gγ while they also lacked plasma membrane distribution (Fig. 1A-Flu). We have previously shown that partial sensitivity to statins is characterized by the complete lack of Gγ at endomembranes, while their presence on the plasma membrane is detectable (13). Interestingly, geranylgeranylated Gγ types (Gγ2, 3, 4, 5, 7, 8, 10, 12, and 13) showed two distinct phenotypes. Gγ2, 3, 4, 7, 8, and 12 showed partial sensitivity, while unexpectedly, the rest Gγ types (Gγ5, 10, and 13) showed a near-complete sensitivity to statins (Fig. 1A-Flu). This raised the question of whether these Gγs (5, 10, and 13) are geranylgeranylated as predicted. To examine their exact type of prenylation, we exposed cells expressing each Gγ subtype to farnesyl transferase inhibitor (FTI), Tipifarnib - 1 µM or geranylgeranyl transferase inhibitor (GGTI), GGTI286 - 10 µM. As expected, Gγ1, 9, and 11 showed a completely inhibited phenotype upon FTI exposure (Fig. 1A-FTI). This also indicated that the remaining Gγ members are geranylgeranylated. Interestingly these FTI insensitive Gγs showed varying sensitivities to GGTI, from moderate to high. Particularly Gγ5, 10 and 13 showed near-complete inhibition while Gγ2, 3, 4, 7, 8 and 12 showed only a partial prenylation inhibition (Fig. 1A-GGTI).

**Figure 1.**
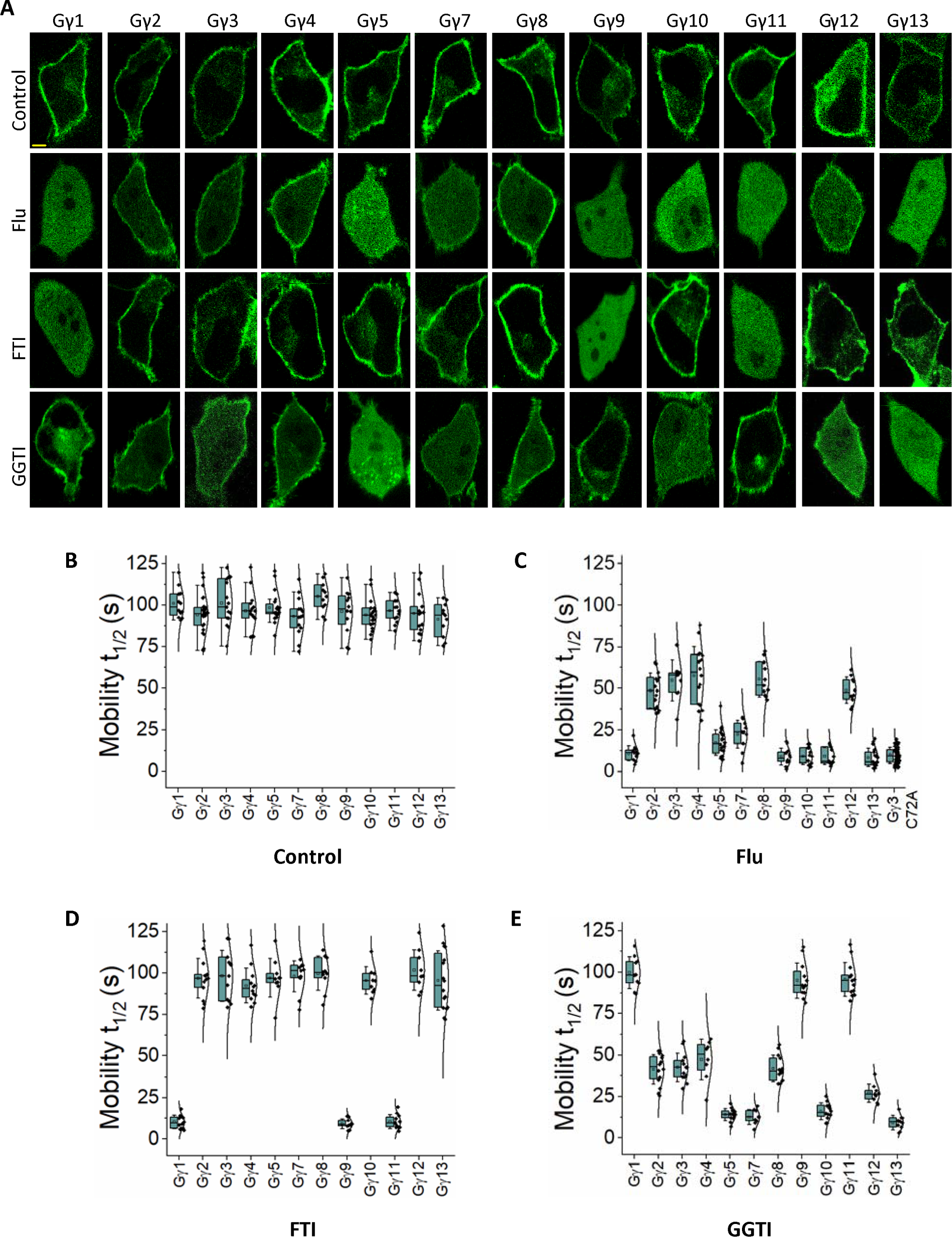
Gγ subtypes show differential sensitivities to statin-induced inhibition of membrane binding. **(A)** Images of HeLa cells expressing YFP tagged Gγ1-5 and 7-13 under the vehicle (control), Fluvastatin (20 µM), FTI (1 µM), or GGTI (10 µM) treated conditions. Images represent the prominent phenotype observed in each population under the given experimental condition (scale: 5 μm; n ≥ 15 for each Gγ type). Whisker box plots generated using fluorescence recovery after half-cell photobleaching of the above cells show the variations in mobility half-time (t_1/2_) of G protein heterotrimers containing different Gγ subtypes, with **(B)** control, **(C)** Fluvastatin-treated, **(D)** FTI-treated, and **(E)** GGTI-treated conditions (Average whisker box plots were plotted using mean±SD; Error bars: SD (Standard deviation); control: n ≥ 12 cells for each Gγ from 154 cells, Fluvastatin-treated: n ≥ 10 cells for each Gγ from 200 cells, FTI-treated: n ≥ 11 cells for each Gγ from 132 cells, GGTI-treated: n ≥ 11 cells for each Gγ 3 from 135 cells (each treatment was performed in 3 ≤ independent experiments; Statistical comparisons were performed using One-way-ANOVA; p<0.05; Flu: Fluvastatin; FTI: Farnesyl transferase inhibitor; GGTI: Geranylgeranyl transferase inhibitor).

Since the visual inspection of Gγ membrane localization inhibition provides only a qualitative estimate of membrane anchorage inhibition, we employed mobilities of Gγ, or lack thereof, as a measure of prenylation. Compared to prenylated Gγ, Gγ with impaired prenylation is expected to have faster mobilities due to reduced membrane interactions (34). To evaluate the mobilities of YFP-tagged Gγ, we examined recovery after photobleaching of half-cell fluorescence and calculated the time to half maximum fluorescence recovery, which we termed mobility half-time (t_1/2_) (Table 1) (35). When proteins are not membrane bound, they move faster since they are cytosolic. However, when only a fraction of the protein population is membrane-anchored, the mobility represents an ensemble movement of both membrane-bound and cytosolic species. Regardless of the Gγ subtype and their type of prenylation, mobility t_1/2_ values of all the control (untreated) Gγs in the heterotrimer were nearly similar, with the average t_1/2_ of 97±4 s (one-way ANOVA: *F*_11,174_ = 1.763, *p* = 0.064) (Fig. 1B). This indicates that Gγs in control cells are in the heterotrimeric form, making their mobility rates Gγ subtype independent (34). When examined, the mobility t_1/2_ of Fluvastatin exposed Gγ1, 9, and 11 were nearly similar and 11±5 s, 9±5 s, and 9±5 s, respectively (one-way ANOVA: *F*_2,33_ = 0.615, *p* = 0.547) (Fig. 1C-Gγ1, 9 and 11). These mobility t_1/2_ values are comparable with the mobility t_1/2_ of the prenylation-deficient Gγ3_C72A_ mutant (10±5 s) that showed a complete cytosolic distribution (one-way ANOVA: *F*_3,84_ = 0.406, *p* = 0.749) (Fig. 1C-Gγ3_C72A_). When Gγ3 expressing cells were exposed to Fluvastatin, the observed mobility t_1/2_ (55±13 s) falls between that of prenylated Gγ3 in control (101±14 s) and the completely cytosolic Gγ3_C72A_ mutant (10±5 s) (Fig. 1C-Gγ3). This mobility rate of Fluvastatin-exposed Gγ3 suggests the presence of both cytosolic and membrane-bound Gγs, indicating partial disruption of membrane anchorage. Compared to untreated conditions, the ∼2-fold reduction in Gγ3 mobility t_1/2_ upon Fluvastatin treatment also signifies the increased cytosolic fraction of Gγ3. Similarly, GGTI exposed Gγ3 also showed an intermediate mobility t_1/2_ (43±9 s) (Fig. 1E-Gγ3), indicating partial geranylgeranylation inhibition. FTI exposed Gγ3 showed a mobility t_1/2_ (98±15 s) that is not significantly different from control Gγ3 (101±14 s) (one-way ANOVA: *F*_1,24_ = 0.250, *p* = 0.621), suggesting that FTI cannot inhibit the activity of geranylgeranyl transferase-I (Fig. 1D-Gγ3). Upon FTI treatment, the mobility t_1/2_ of farnesylation-sensitive Gγs (Gγ1, 9, and 11) are significantly reduced (∼9-10 s) while that of geranylgeranylation-sensitive Gγs remained nearly unchanged, confirming the exclusive farnesylation-sensitivities of these Gγs (Fig. 1D). Nevertheless, upon exposing cells to GGTI, only geranylgeranylation-sensitive Gγs showed a significant reduction in mobility t_1/2_ (Fig. 1E), while their mobility was unaffected upon FTI exposure (Fig. 1D). However, GGTI-induced prenylation inhibition sensitivity of geranylgeranylation-sensitive Gγs also showed a wide range from near complete (as in Gγ5, 10, and 13) to partial (as in Gγ2, 3, 4, 7, 8, and 12).

**Table 1:** Mobility properties of Gγ types.

| Gγ type | Ct Sequence | Mobility $t_{1/2}$ (s) | | | |
| --- | --- | --- | --- | --- | --- |
|  |  | Control | Flu* | FTI* | GGTI* |
| Gγ1-WT | N P F K E L K G G C V I S | 101 ± 9 | 11 ± 5 | 10 ± 4 | 100 ± 10 |
| Gγ2-WT | N P F R E K K F F C A I L | 90 ± 11 | 49 ± 10 | 97 ± 12 | 41 ± 9 |
| Gγ3-WT | N P F R E K K F F C A I L | 102 ± 14 | 55 ± 12 | 98 ± 15 | 43 ± 9 |
| Gγ4-WT | N P F R E K K F F C T I L | 97 ± 11 | 58 ± 17 | 92 ± 10 | 47 ± 12 |
| Gγ5-WT | N P F R P Q K V - C S F L | 101 ± 9 | 17 ± 8 | 97 ± 12 | 14 ± 4 |
| Gγ7-WT | N P F K D K K P - C I I L | 94 ± 12 | 22 ± 8 | 98 ± 9 | 13 ± 4 |
| Gγ8-WT | N P F R D K R L F C V I L | 105 ± 9 | 55 ± 11 | 100 ± 10 | 42 ± 8 |
| Gγ9-WT | N P F K E - K G G C I I S | 96 ± 13 | 9 ± 5 | 9 ± 3 | 95 ± 10 |
| Gγ10-WT | N P F R E P R S - C A I L | 97 ± 9 | 9 ± 5 | 95 ± 8 | 16 ± 5 |
| Gγ11-WT | N P F K E - K G S C V I S | 97 ± 7 | 9 ± 5 | 10 ± 4 | 96 ± 10 |
| Gγ12-WT | N P F K D K K T - C I I L | 96 ± 7 | 49 ± 8 | 102 ± 12 | 27 ± 5 |
| Gγ13-WT | N P W V E K G K - C T I L | 95 ± 12 | 8 ± 5 | 95 ± 18 | 9 ± 4 |

\*Mobility $t_{1/2}$ of major phenotype

### 3.2 Statin-induced Gγ delocalization to the cytosol is prenylation-type independent

Our previous work has established that Gγ subunits such as γ9, γ1, and γ11 show faster translocation with t_1/2_ of a few seconds and greater magnitudes. They were identified as low membrane affinity Gγs. Conversely, Gγ with larger translocation t_1/2_ values and lower magnitudes, including Gγ2, γ3, and γ4, are considered to be high membrane affinity Gγs (40). Interestingly, when examined, the fastest translocating Gγ9 and the slowest translocating Gγ3 also exhibited two distinct sensitivities to Fluvastatin-induced membrane anchorage inhibition; Gγ9 with near-complete cytosolic distribution and Gγ3 with partial cytosolic distribution (Fig. 1A and C) (13). To quantitatively define these two classifications (complete cytosolic vs partial cytosolic), we assigned Gγs with 0-24 s mobility t_1/2_ values as complete cytosolic and Gγ with 25-60 s mobility t_1/2_ as partial cytosolic. According to this classification, upon Fluvastatin exposure, ∼100% of the Gγ9 cells showed a prominently increased cytosolic fluorescence due to cytosolic Gγ, while the fraction of cells with partial cytosolic distribution was nearly zero and insignificant (Fig. 2A, 2B and Table 3, Gγ9-WT). Interestingly, ∼99% of Gγ3 cells showed partial inhibition of membrane binding while near-complete inhibition was inconsequential with Fluvastatin treatment (Fig. 2A, 2B and Table 3, Gγ3-WT). To examine the contribution of the CaaX sequence in determining the degree of statin-induced retardation of membrane binding, we compared CaaX motif mutants of Gγ9 and Gγ3 with their corresponding wild types. Similar to Gγ9-WT, when exposed to Fluvastatin, ∼99% of cells expressing a Gγ9 mutant with the CaaX sequence of Gγ3 (Gγ9_CALL_) exhibited near-complete cytosolic distribution indicating complete disruption of membrane binding (Fig. 2A, 2B and Table 3, Gγ9_CALL_). However, the FTI and GGTI sensitivities of the Gγ9_CALL_ mutant showed that it is geranylgeranylated as predicted from its CaaX (CALL) sequence (Fig. 2A, Gγ9_CALL_). Similarly, the Gγ3 mutant containing the CaaX sequence of Gγ9 (Gγ3_CIIS_) showed a partial disruption of membrane anchoring when cells were exposed to Fluvastatin even though it is farnesylation sensitive (Fig. 2A, Gγ3_CIIS_). This response was prominent in ∼97% of the mutant-expressing cells, while only ∼3% of the total population exhibited near-complete inhibition of membrane binding (Fig. 2B and Table 3-Gγ3_CIIS_). We also calculated mobility rates of the wild type and two Gγ mutants (Fig. 2C and Table 2). Even though switching the CaaX motif convincingly switched the FTI and GGTI sensitivities of Gγ9 and Gγ3, the Fluvastatin sensitivity remained unchanged between the WT controls and the corresponding mutants, implying distinct molecular determinants governing the statin sensitivity of Gγ.

**Figure 2.**
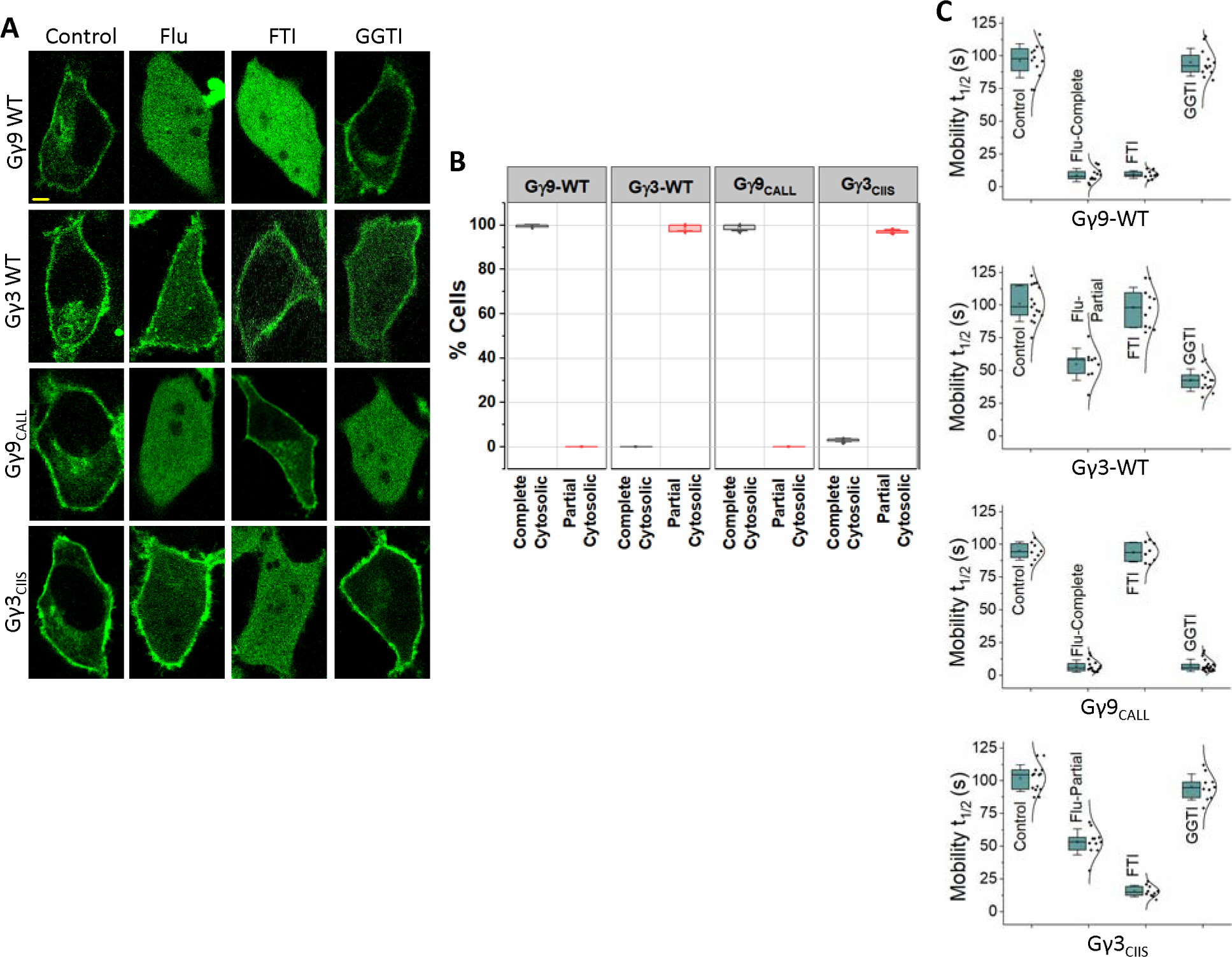
Statin sensitivity and prenylation efficacy of Gγ are prenylation type independent. **(A)** Subcellular distribution of GFP tagged wild type Gγ9, Gγ3, and mutants Gγ9_CALL_, and Gγ3_CIIS_, with vehicle (control), Fluvastatin (20 µM), FTI (1 µM), or GGTI (10 µM) treated conditions. Images represent the prominent phenotype observed in each population under the given experimental condition (scale: 5 μm; n ≥ 15 for each Gγ type). **(B)** Grouped box chart shows the percentages of cells in each Gγ type (Gγ9 WT, Gγ3 WT, Gγ9_CALL_, and Gγ3_CIIS_) showing near-complete (black) and partial (red) cytosolic distribution with Fluvastatin treatment. **(C)** The whisker box plots show mobility half-time (t_1/2_) of proteins in (A) determined using fluorescence recovery after half-cell photobleaching. (Average box plots were plotted using mean±SD; Error bars: SD (Standard deviation); Gγ9 WT: n= 571 total number of cells from 7 independent experiments, Gγ3 WT: n= 597 total number of cells from 7 independent experiments, Gγ9_CALL_: n= 624 total number of cells from 7 independent experiments, Gγ3_CIIS_: n=408 total number of cells from 5 independent experiments; Statistical comparisons were performed using One-way-ANOVA; p<0.05; Flu: Fluvastatin; FTI: Farnesyl transferase inhibitor; GGTI: Geranylgeranyl transferase inhibitor; WT: wild-type).

**Table 2:**
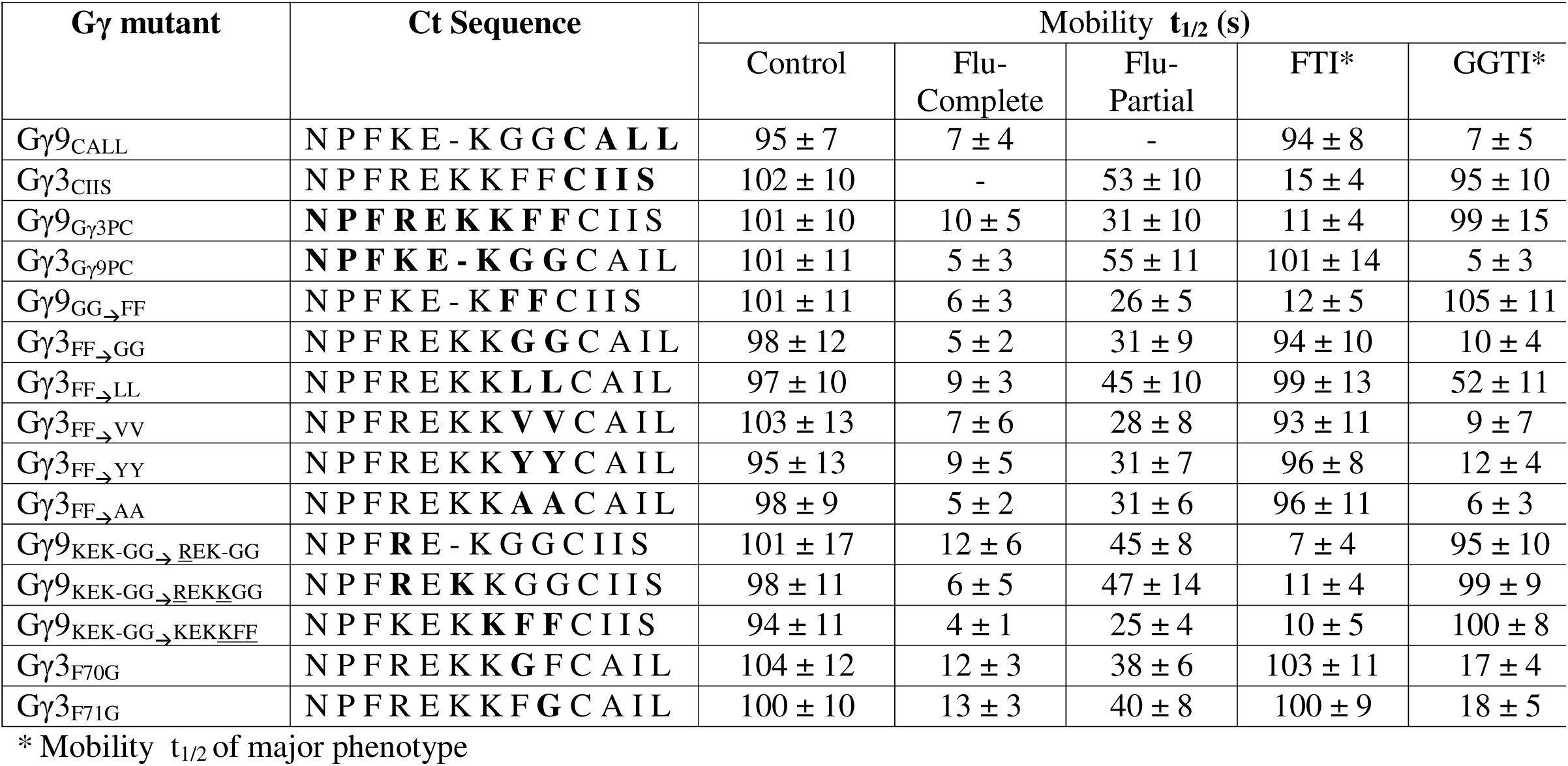
Mobility properties of Gγ mutants.

**Table 3:** Fluvastatin-induced prenylation inhibition (or partial inhibition) of Gγ3, Gγ9 and their Ct mutants.

| Gγ mutant | Ct Sequence | Percent cells (%) | |
| --- | --- | --- | --- |
|  |  | Complete | Partial |
| Gγ9-WT | N P F K E - K G G C I I S | 99.7 ± 0.5 | 0 |
| Gγ3-WT | N P F R E K K F F C A I L | 0 | 98.7 ± 1.2 |
| Gγ9<sup>CALL</sup> | N P F K E - K G G C A L L | 98.6 ± 1.2 | 0 |
| Gγ3<sup>CIIS</sup> | N P F R E K K F F C I I S | 2.9 ± 0.7 | 97.1 ± 0.7 |
| Gγ9<sup>Gγ3PC</sup> | N P F R E K K F F C I I S | 23.8 ± 12.9 | 74.5 ± 13.4 |
| Gγ3<sup>Gγ9PC</sup> | N P F K E - K G G C A I L | 99.3 ± 0.9 | 0 |
| Gγ9<sup>GG→FF</sup> | N P F K E - K F F C I I S | 87.4 ± 2.7 | 12.5 ± 2.7 |
| Gγ3<sup>FF→GG</sup> | N P F R E K K G G C A I L | 94.5 ± 1.3 | 5.5 ± 1.3 |
| Gγ3<sup>FF→LL</sup> | N P F R E K K L L C A I L | 4.1 ± 3.1 | 95.9 ± 3.1 |
| Gγ3<sup>FF→VV</sup> | N P F R E K K V V C A I L | 88.2 ± 4.5 | 11.8 ± 4.5 |
| Gγ3<sup>FF→YY</sup> | N P F R E K K Y Y C A I L | 87.9 ± 2.3 | 12.1 ± 2.3 |
| Gγ3<sup>FF→AA</sup> | N P F R E K K A A C A I L | 92.8 ± 3.0 | 7.2 ± 3.0 |
| Gγ9<sup>KEK-GG→REK-GG</sup> | N P F R E - K G G C I I S | 83.6 ± 1.6 | 15.7 ± 0.7 |
| Gγ9<sup>KEK-GG→REKKGG</sup> | N P F R E K K G G C I I S | 65.2 ± 3.8 | 35.7 ± 5.9 |
| Gγ9<sup>KEK-GG→KEKKFF</sup> | N P F K E K K F F C I I S | 51.3 ± 6.3 | 48.5 ± 6.4 |
| Gγ3<sup>F70G</sup> | N P F R E K K G F C A I L | 28.9 ± 1.1 | 72.6 ± 0.9 |
| Gγ3<sup>F71G</sup> | N P F R E K K F G C A I L | 20.0 ± 0.4 | 77.7 ± 5.6 |

### 3.3 Pre-CaaX sequences of Gγs control their statin sensitivity

Since Fluvastatin-induced membrane binding inhibition profiles of Gγ family members were Gγ-type dependent, we examined the molecular reasoning for this behavior, especially considering that Fluvastatin sensitivity of Gγ is independent of their prenylation type. We have extensively documented that, in addition to prenyl and carboxymethyl modifications on the Ct *Cys* of Gγ, its adjacent pre-CaaX region also controls Gβγ-membrane interactions (40–42). Our previous work showed the crucial involvement of pre-CaaX residues in specific Gγs regulating their membrane affinity (41). Therefore, we examined whether the chemistry of pre-CaaX amino acids also controls the protein’s statin sensitivity (Fig. 3A). For this, we computationally determined the hydrophobicity of the Ct polypeptide of Gγ only comprising pre-CaaX and CaaX regions (before prenylation), using octanol-water partition coefficient (*K_OW_*)-based *Log Cavity Energy* (*Log CE*) calculation (please see Methods 2.5) (Fig. 3B, Table S1). It has been shown that the difference between the free energy of cavity formation in the organic and water solvents indicates a peptide’s relative affinity between the two phases and its hydrophobicity (43). All the Gγ-derived peptides showed higher positive ΔG values for water (ΔG_W_). Interestingly, Gγ2, 3, 4, 5, and 8 exhibited lower but positive ΔG values for octanol (ΔG_O_), indicating that they have a higher affinity to the lipid phase (Table S1).

**Figure 3.**
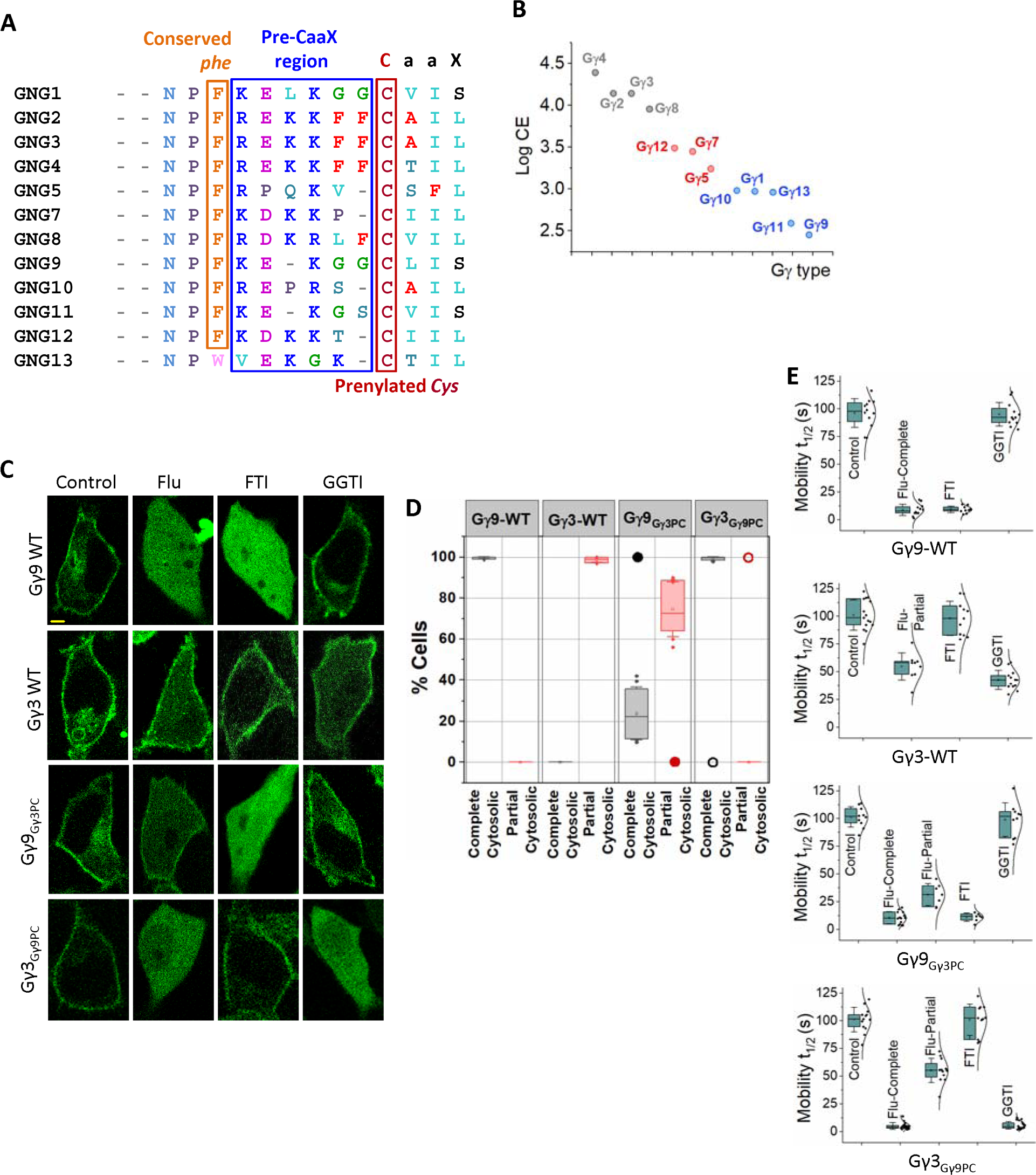
The pre-prenylation sequence determines statin sensitivity and prenylation efficacy of Gγ. **(A)** Comparison of Ct domain amino acid sequences of Gγ. The Ct domain of Gγ consists of the starting conserved NPF sequence, followed by the middle pre-CaaX region and the final CaaX motif. Gγ subtypes with a C-terminal *Ser* (Balck) at the ‘X’ position are farnesylated. Gγs containing *Leu* (light blue) at the ‘X’ position of CaaX are geranylgeranylated. During prenylation, the prenyl moiety is attached to the prenylated *Cys* (maroon), and this *Cys* is further modified during post-prenylation processing (proteolysis at the C-terminal three -aaX residues by the Ras converting CaaX endopeptidase 1 (RCE1), and then carboxymethylation of the new isoprenylcysteine C-terminus by isoprenylcysteine carboxyl methyltransferase (ICMT)). In the pre-CaaX region, hydrophobic *Phe* residues are shown in red, and positively charged residues are in blue. **(B)** The scatter plot shows the log CE values to measure the hydrophobicity of pre-CaaX+CaaX Ct polypeptide regions of Gγ. **(C)** Images of HeLa cells expressing GFP tagged Gγ9-WT, Gγ3-WT, Gγ9_Gγ3PC_, and Gγ3_Gγ9PC_, exposed to vehicle (Control), Fluvastatin (20 µM), FTI (1 µM), or GGTI (10 µM). Images represent the prominent phenotype observed in each population under given experimental conditions (scale: 5 μm; n ≥ 15 for each Gγ type). **(D)** Grouped box chart shows the percentages of cells in each Gγ mutant (Gγ9_Gγ3PC_, Gγ3_Gγ9PC_, Gγ9_GG→FF_, and Gγ3_FF→GG_ mutants) showing near-complete (black) and partial (red) cytosolic distribution with Fluvastatin treatment. The black/red circles indicate the % cells showed near-complete or partial cytosolic distribution in each corresponding wild-type Gγ expressing cells. **(E)** The whisker box plots show mobility half-time (t_1/2_) of proteins in (C) determined using fluorescence recovery after half-cell photobleaching. (Average box plots were plotted using mean±SD; Error bars: SD (Standard deviation); Gγ9 WT: n= 571 total number of cells from 7 independent experiments, Gγ3 WT: n= 597 total number of cells from 7 independent experiments, Gγ9_Gγ3PC_: n= 686 total number of cells from 8 independent experiments, Gγ3_Gγ9PC_: n= 754 total number of cells from 8 independent experiments; Statistical comparisons were performed using One-way-ANOVA; p < 0.05; Flu: Fluvastatin; FTI: Farnesyl transferase inhibitor; GGTI: Geranylgeranyl transferase inhibitor; PC: pre-CaaX).

The cavity energy of pre-CaaX+CaaX peptides of Gγ indicates that Gγ4, 3, 2, and 8 possess significantly higher stability in lipids compared to Gγ1, 9, and 11. Interestingly, log CE of pre-CaaX+CaaX of Gγ10 and 13 also showed less favorable values for lipids in the same range as Gγ1, 9, and 11 (Fig. 3B, Table S1). These log CE values agree with our unexpected observation that similar to Gγ1, 9, 11, geranylgeranylated Gγ10, 13 also possess similar sensitivity to Fluvastatin (Fig. 1 and 3B). This aligns with our hypothesis that unprenylated polypeptide regulates the prenylation efficacy by interacting differentially with membranes where prenylation occurs. Further, the pre-CaaX+CaaX peptides from Gγ5, 7, and 12 that showed log CE values between the above two groups (Gγ2, 3, 4, 8 vs. Gγ1, 9, 11), also exhibited partial, however, more significant prenylation inhibition than Gγ2, 3, 4, 8 upon Fluvastatin, supporting our hypothesis (Fig. 3B and Table S1).

Since these findings suggest the involvement of the pre-CaaX region in regulating Gγ-membrane interactions, and different Gγ types show distinct sensitivities to statin, regardless of the type of prenylation, we tested the hypothesis that the pre-CaaX region hydrophobicity of Gγ polypeptide govern their statin sensitivity. We first examined the influence of exchanging pre-CaaX sequences between Gγ3 and Gγ9, representative Gγs of the two extremes of Gγ membrane affinities (40). We comparatively examined the statin sensitivity of the Gγ9 mutant containing the pre-CaaX (PC) of Gγ3 (Gγ9_Gγ3PC_ or -**<u>NPFREKKFF</u>**CLIS) that we reported previously (41) and a new Gγ3 mutant containing the pre-CaaX of Gγ9 (Gγ3_Gγ9PC_ or -**<u>NPFKEKGG</u>**CALL). Interestingly, in the presence of Fluvastatin, cells expressing Gγ9_Gγ3PC_ exhibited a prominent partial inhibition of membrane binding (note the near-complete cytosolic distribution in WT Gγ9-black filled circle in Fig. 3D), while Gγ3_Gγ9PC_ showed a near-complete inhibition of membrane anchorage (compare the partial cytosolic distribution in WT Gγ3-Red open circle in Fig. 3D) (Fig. 3C, 3D and Table 3-Gγ9_Gγ3PC_ and Gγ3_Gγ9PC_). Though the images here represent the most abundant phenotype, the grouped box chart shows the percent abundance of each phenotype of the respective Gγ mutant (Fig. 3C and 3D). Interestingly, compared to Gγ9-WT, Gγ9_Gγ3PC_ also shows a significant reduction in the near-complete inhibition phenotype (from ∼100% in WT to ∼24% in the mutant) (Fig. 3C, 3D and Table 3-Gγ9_Gγ3PC_, black). This phenotype additionally showed a significantly faster mobility compared to that of the partial cytosolic phenotype (complete cytosolic: 5±3 s, partial cytosolic: 55±11 s, one-way ANOVA: *F*_1,36_ = 477.158, *p* = 0.0001) (Fig. 3E and Table 2-Gγ9_Gγ3PC_), which is the most abundant phenotype (∼75%) of Fluvastatin-exposed cells (Fig. 3D - Gγ9_Gγ3PC_, red). However, Fluvastatin exposed Gγ3_Gγ9PC_ cells only exhibited near-complete cytosolic phenotype (from 0% in WT to ∼99% in the mutant) (Fig. 3D and Table 3-Gγ3_Gγ9PC_, black), eliminating the partial cytosolic phenotype (Fig. 3D and Table 3-Gγ3_Gγ9PC_, red). Mobility data confirmed the complete inhibition of membrane association (Fig 3E and Table 2-Gγ3_Gγ9PC_). This is distinctly different from the response of WT Gγ3 cells to Fluvastatin treatment (Fig. 2B, Fig. 3D-Red open circle). Further, by exposing these pre-CaaX mutants to prenyltransferase inhibitors, we show that pre-CaaX switching did not change their type of prenylation (Fig. 3C-FTI and GGTI-bottom two panels). For instance, similar to Gγ9-WT, the Gγ9_Gγ3PC_ mutant showed sensitivity to FTI but not to GGTI. We have confirmed this phenotype data by examining their mobility t_1/2_ (Fig. 3E and Table 2-Gγ9_Gγ3PC_ and Gγ3_Gγ9PC_).

### 3.4 Hydrophobicity character of near prenyl-Cys residues indicates statin sensitivity of Gγ

Next, we focused on the pre-CaaX region hydrophobicity in determining the statin sensitivity of Gγ. We employed a Gγ3 mutant, in which the *Phe-duo* adjacent to prenyl-*Cys* is replaced with two *Gly* residues (Gγ3_FF→GG_: NPFREKK**<u>GG</u>**CALL), and a Gγ9 mutant carrying two *Phe* residues in place of *Gly* (Gγ9_GG→FF_: NPFKEK-**<u>FF</u>**CLIS) (41). Unlike in Gγ9-WT, which showed ∼100% near-complete cytosolic phenotype, the Gγ9_GG→FF_ mutant showed a significant reduction in membrane anchorage inhibition down to ∼69% while its partial inhibition cytosolic population increased to ∼31% (Fig. 4B, C and Table 3-Gγ9_GG→FF_). These full and partial phenotypes were further confirmed using mobility rates (Fig. 4D and Table 2-Gγ9_GG→FF_). However, the anchorage inhibitory effect upon FTI, but not due to GGTI, showed that the prenylation type of this Gγ9_GG→FF_ mutant remained unchanged (farnesylated). Similarly, compared to Gγ3-WT, however, more substantially, the Fluvastatin exposed Gγ3_FF→GG_ mutant cells exhibited a higher susceptibility to membrane anchorage inhibition, in which the near-complete cytosolic distribution became the prominent phenotype (∼83%) (Fig. 4B, C and Table 3-Gγ3_FF→GG_, black and 4D and Table 2-Gγ3_FF→GG_), while the partial inhibition was reduced (∼17%) (Fig. 4B-D and Table 3-Gγ3_FF→GG_, red). The sensitivity to GGTIs and lack thereof to FTIs indicated that the type of prenylation of this Gγ3_FF→GG_ mutant remained unchanged (geranylgeranylated) (Fig. 4B and D-Gγ3_FF→GG_). Building on these observations, we propose that Gγ9_GG→FF_ is a gain-of-function mutant in which the gain is the resistance to statin sensitivity.

**Figure 4.**
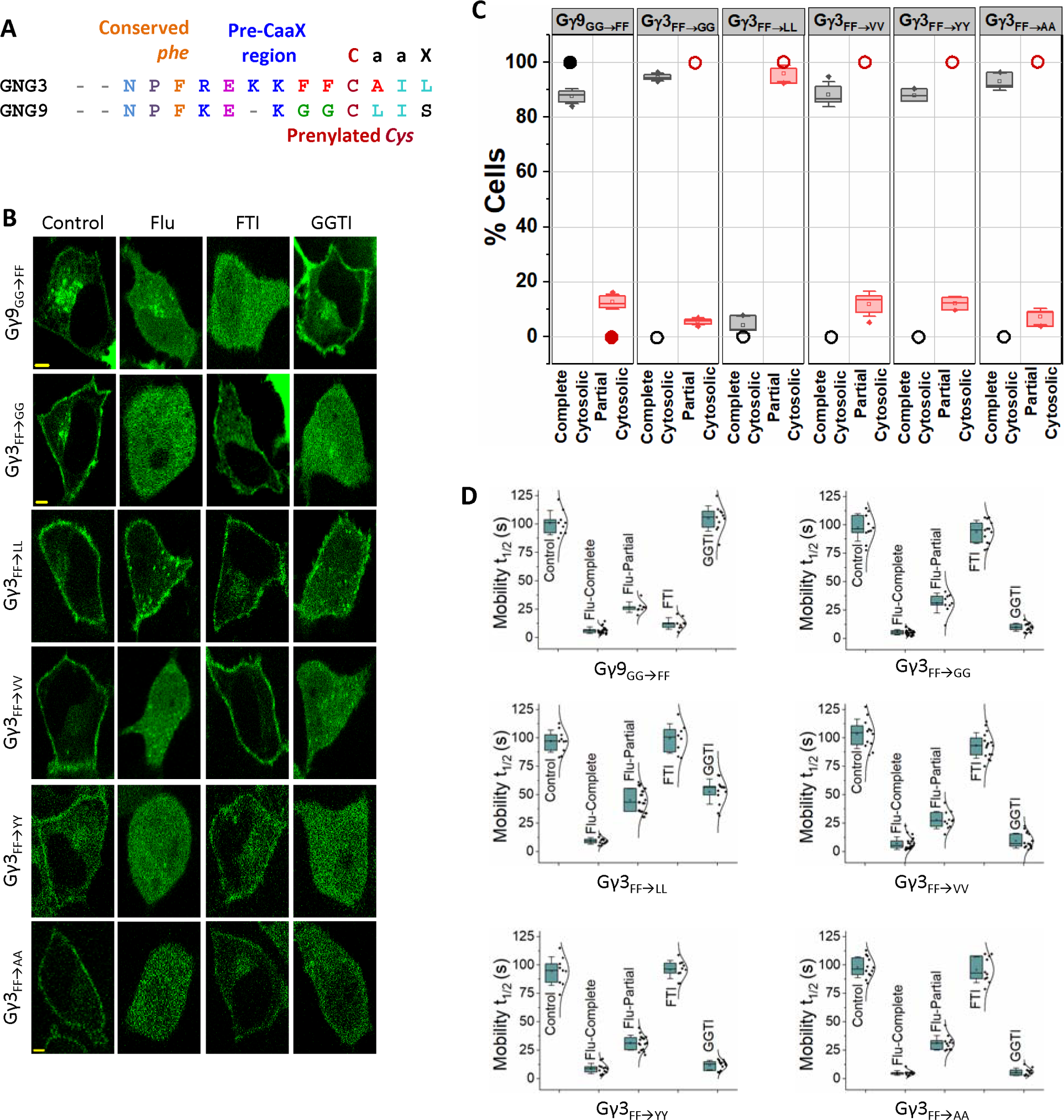
Hydrophobic residues adjacent to prenylated *Cys* contribute significantly to the prenylation efficacy and statin sensitivity of Gγ. **(A)** Comparison of Ct domain amino acid sequences of Gγ3-WT and Gγ9-WT. **(B)** Images of HeLa cells expressing GFP tagged Gγ9_GG→FF_, Gγ3_FF→GG_, Gγ3_FF→LL_, Gγ3_FF→VV_, Gγ3_FF→YY_, and Gγ3_FF→AA_ mutants exposed to vehicle (Control), Fluvastatin (20 µM), FTI (1 µM), or GGTI (10 µM). Images represent the prominent phenotype observed in each population under given experimental conditions (scale: 5 μm; n ≥ 15 for each Gγ type). **(C)** Grouped box chart shows the percentages of cells in each Gγ mutant (Gγ9_GG→FF_, Gγ3_FF→GG_, Gγ3_FF→LL_, Gγ3_FF→VV_, Gγ3_FF→YY_, and Gγ3_FF→AA_) showing near-complete (black) and partial (red) cytosolic distribution with Fluvastatin treatment. The black/red open circles indicate the % cells with near-complete or partial cytosolic distribution in each corresponding wild-type Gγ (Gγ3-WT) expressing cells. **(D)** The whisker box plots show mobility half-time (t_1/2_) of proteins in (B) determined using fluorescence recovery after half-cell photobleaching. (Average box plots were plotted using mean±SD; Error bars: SD (Standard deviation); Gγ9_GG→FF_: n= 884 total number of cells from 9 independent experiments, Gγ3_FF→GG_: n= 225 total number of cells from 3 independent experiments, Gγ3_FF→LL_: n= 428 total number of cells, Gγ3_FF→VV_: n= 364 total number of cells, Gγ3_FF→YY_: n= 407 total number of cells, Gγ3_FF→AA_: n= 358 total number of cells; Each Gγ mutant was examined in 3 independent experiments; Statistical comparisons were performed using One-way-ANOVA; p < 0.05; Flu: Fluvastatin; FTI: Farnesyl transferase inhibitor; GGTI: Geranylgeranyl transferase inhibitor; WT: wild-type).

However, the gain here is smaller than the significant loss observed in Gγ3_FF→GG_ that we identified as the loss of function mutant. We then examined the individual contribution of each *Phe* residue in the *Phe-duo* towards prenylation efficacy. We generated two Gγ3 mutants, Gγ3_F70G_ and Gγ3_F71G,_ and examined their statin sensitivity (Fig. S1). Based on their sensitivity to GGTI, we confirmed that both mutants are geranylgeranylation sensitive (Fig. S1). Compared to Gγ3-WT (Fig. 1B and 1C), a significantly higher fraction of both mutant cell populations showed near-complete inhibition of membrane localization (Gγ3_F70G_: ∼29%, Gγ3_F71G_: ∼20%) (Fig. S1 and Table 3-Gγ3_F70G_ and Gγ3_F71G_-black). Considering that the near-complete inhibition is absent in Gγ3-WT, these data indicate a crucial role of each *Phe* for Gγ3 to gain its efficient prenylation. Further, compared to the 55±13 s mobility t_1/2_ observed in Fluvastatin exposed Gγ3-WT, both the mutants showed enhanced mobility (Gγ3_F70G_: 38±6 s, Gγ3_F71G_: 40±8 s) in their partial inhibition populations (Table 2). This suggests that the extent of statin sensitivity in the partially inhibited population is much greater in the absence of each residue of the *Phe-duo*. These data collectively demonstrate that the hydrophobic character of the residues adjacent to prenyl-*Cys* is a crucial determinant of G proteins’ statin sensitivity.

To further confirm the influence of prenyl-*Cys* adjacent hydrophobic residues on the protein’s statin sensitivity, we altered the Gγ pre-CaaX hydrophobicity by mutating the *Phe-duo* in Gγ3. When the *Phe-duo* is mutated to two *Leu* residues, which has a similar hydrophobicity as *Phe* (44, 45), similar to the wild-type (Gγ3 WT), the majority of the Gγ3_FF LL_ (NPFREKK**<u>LL</u>**CALL) mutant cells still showed partial inhibition of membrane anchorage (∼96%) primarily (Fig. 4B, C and Table 3-Gγ3_FF LL_, red), while a minor percentage (∼4%) showed a near-complete inhibition (Fig. 4C and Table 3-Gγ3_FF LL_, black). Interestingly mutants generated by replacing the *Phe-duo* either with *Val*, *Ala*, or *Tyr* (Gγ3_FF→VV_, Gγ3_FF→AA_, or Gγ3_FF YY_), which possess significantly lower hydrophobicity indices than *Phe* (44, 45), showed a prominent (∼95-100%) near-complete membrane localization inhibition upon Fluvastatin exposure (Fig. 4B, C and Table 3). To obtain quantitative data corroborating the imaging observations, we next examined mobilities of mutant Gγ subunits under pharmacological perturbation. While the mobility of partially inhibited Gγ3_FF LL_ was similar to that of Gγ3-WT (Fig. 1C-Gγ3 WT), completely inhibited populations of the rest of the mutants containing less hydrophobic residues adjacent to their prenyl-*Cys* (Gγ3_FF→VV_, Gγ3_FF→YY_, Gγ3_FF→AA_) exhibited significantly faster mobility upon Fluvastatin treatment (Fig. 4D and Table 2). The similar and substantially slower mobility of Fluvastatin-treated Gγ3-WT and Gγ3_FF LL_ indicates the considerable presence of a prenylated Gγ population bound to membranes, suggesting their significant resistance to membrane localization inhibition by statins. Contrastingly, significantly faster mobility of Gγ3_FF→VV_, Gγ3_FF→YY_, and Gγ3_FF→AA_ (Fig. 4D and Table 2) were also closer to the mobility of cytosolic Gγ9-WT observed in Fluvastatin-treated cells (Fig. 1C-Gγ9 WT), indicating that even for G proteins undergoing geranylgeranylation, the hydrophobic character of the pre-CaaX is a crucial regulator of their prenylation process. We also observed that the above Gγ3 mutations did not alter the prenylation type (Fig. 4B, Dand Table 2, FTI and GGTI).

Interestingly, our data show that GGTI-mediated geranylgeranylation inhibition is partial in highly hydrophobic pre-CaaX residues carrying Gγ3 WT and Gγ3_FF LL_ mutant cells. On the contrary, GGTI induced a highly effective near-complete inhibition of geranylgeranylation in Gγ3_FF→VV_, Gγ3_FF→AA_, or Gγ3_FF→YY_ mutants (Fig. 4B and D). Here, we hypothesize that compared to the above three mutants, the observed partial prenylation in GGTI-exposed Gγ3-WT and Gγ3_FF LL_ mutant cells results from the enhanced hydrophobicity of their pre-CaaX region. This may allow these Gγs to undergo geranylgeranylation to a significant extent, likely using the residual geranylgeranyl transferase activity remaining in the cell. Further supporting the significance of pre-CaaX residues in regulating the prenylation process, mutant cells exposed to GGTI also showed mobility half times similar to the values observed in cells exposed to Fluvastatin (Fig. 4D and Table 2). Therefore, these data suggest that the hydrophobicity of the pre-CaaX region is a primary regulator of a G protein’s resistivity to prenyltransferase inhibitors.

### 3.5 Optogenetically-controlled reversible unmasking-masking confirms the significance of Ct hydrophobic residues on prenylation and the membrane affinity of the prenylated protein

To examine how the prenylated-*Cys*-adjacent hydrophobic residues first influence prenylation and then regulate membrane binding of the prenylated protein, we engineered an improved light-induced dimer (iLID) system-based, blue-light-gated, monomeric photo-switch to expose and mask the *Phe-duo* of a protein with a geranylgeranylating CALL at the Ct (iLID_FCALL_). Since iLID is ending with a *Phe* (F449), the Ct sequence of the protein is — FFCALL (46) (Fig. 5A-top and B). Upon prenylation, this protein should become iLID_F_ with ending geranylgeranylated and carboxymethylated *Cys.* With the incorporated second *Phe* (F550), we created a *Phe-duo* (Fig. 5A-top) in the protein Venus-iLID_FCALL_ (Fig. 5B). Upon blue light exposure, both before and after prenylation, we expected the *Phe-duo* of the protein to be exposed (Fig. 5B).

**Figure 5.**
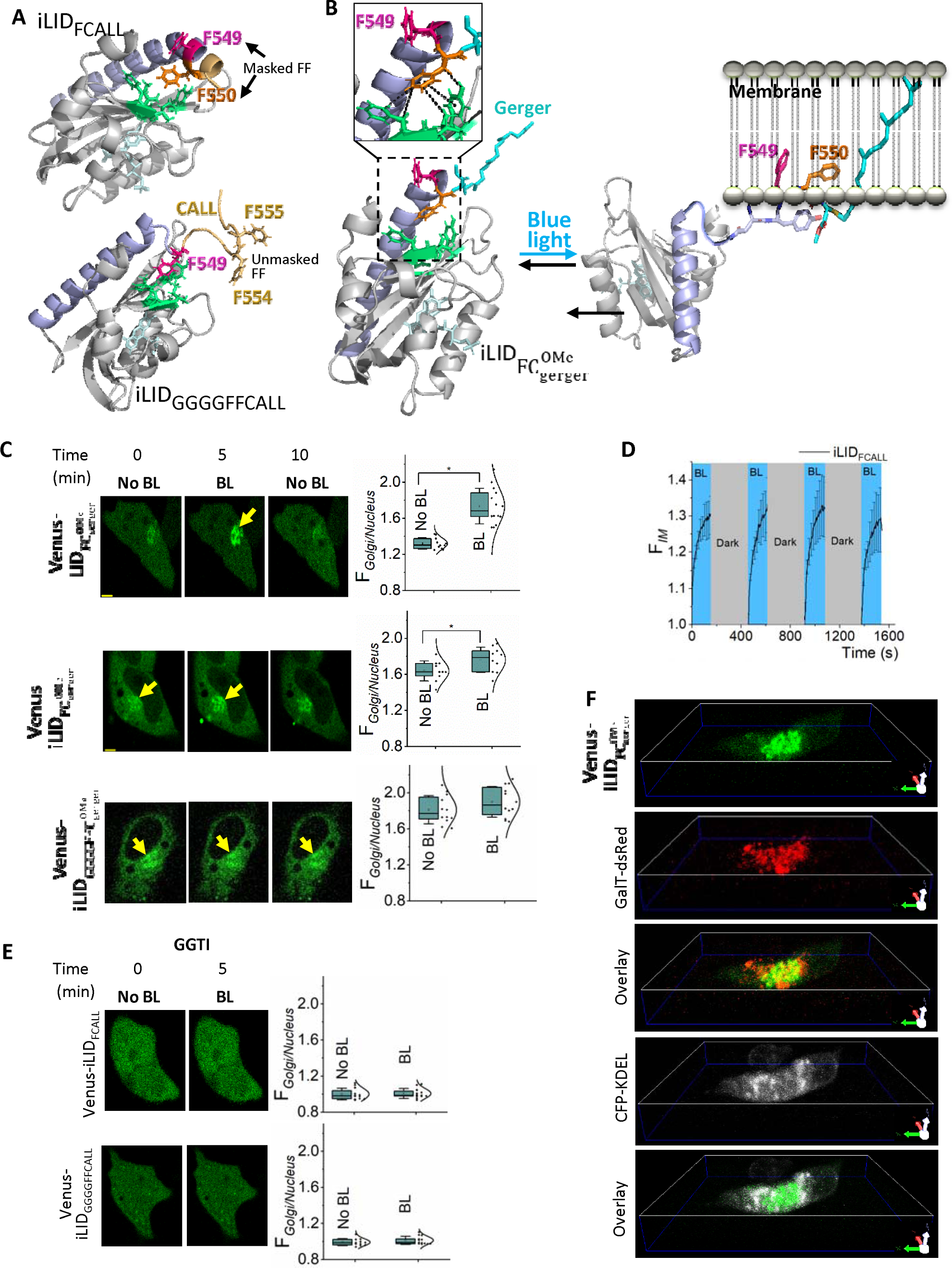
Optogenetic amino acid masking-unmasking shows pre-CaaX hydrophobicity-dependent prenylation efficacy and post-prenylation behavior regulation of proteins. **(A)** The modeled structures of the two photoswitches, iLID_FCALL_ and iLID_GGGGFFCALL_ (based on PDB ID-4WF0) before their prenylation. Resembling Gγ3 *Phe-duo*, the introduced *Phe550* (orange) in iLID_FCALL_ (Top) creates the *Phe-duo* using *Phe549* (pink) of iLID. As a control, an iLID variant with an exposed *Phe-duo* (*Phe554* and *Phe555-*light orange) was generated by placing a linker sequence (GGGG-light orange loop region) between iLID and *Phe-duo* (FF) (iLID_GGGGFFCALL_-Bottom). The introduced F550 in iLID_FCALL_ fits into a hydrophobic pocket on the surface of the per-arnt-sim (PAS) domain made up of I417, I428, F429, and Y508 (green), while in iLID_GGGGFFCALL_ F549 fits into this hydrophobic pocket allowing the *Phe-duo* to be exposed**. (B)** Optogenetic regulation of prenylated 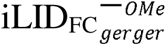. Blue light-induced Jα helix relaxation results in iLID’s SsrA peptide unmasking, exposing the *Phe-duo,* likely promoting membrane interaction of iLID_F_ by inserting the *Phe-duo* (F549 and F550) and the geranylgeranyl moiety into the hydrophobic tail region of the lipid bilayer. The magnified view shows the hydrophobic interactions of the F549 in 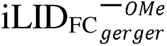 with the hydrophobic pocket of the PAS domain before blue light irradiation. **(C)** Images of HeLa cells expressing Venus-iLID_FCALL_, overnight blue light exposed Venus-iLID_FCALL_ or Venus-iLID_GGGGFFCALL_ (control) show the initial subcellular distribution of the proteins and their blue light-induced changes. In the control protein, the yellow arrow indicates the Golgi/endomembrane distribution of the constructs (Scale bar: 5 µm). The whisker box plots show the Golgi:Nucleus Venus fluorescence ratio under each condition (Average box plots were plotted using mean±SD; Error bars: SD (Standard deviation); Statistical comparisons were performed using One-way-ANOVA; p < 0.05) **(D)** The plot shows the reversible recruitment of iLID_FCALL_ to the Golgi upon blue light (error bars: SD, 10 < n). **(E)** Images of HeLa cells expressing Venus-iLID_FCALL_ or Venus-iLID_GGGGFFCALL_ show a completely cytosolic distribution of the two proteins when the cells are exposed to GGTI, before and after blue light irradiation (10 µm). **(F)** 3-D images of blue light irradiated HeLa cells expressing Venus-iLID_FCALL_, GalT-dsRed, and CFP-KDEL. Blue light-induced Golgi recruitment of Venus-iLID_FCALL_ is confirmed by its colocalization with the Golgi marker GalT-dsRed and non-overlapping distribution with the ER marker CFP-KDEL (Scale bar: 5 µm) (BL: Blue light; O/N BL: Overnight blue light; GGTI: Geranylgeranyl transferase inhibitor).

When Venus-iLID_FCALL_ was expressed in HeLa cells, it showed primarily a cytosolic distribution with a minor Golgi localization (Fig. 5C-top, No BL). To quantify the subcellular distribution of this protein, we calculated the normalized Venus fluorescence ratio of the Golgi and nucleus. Since the prenylation-lacking Venus-iLID_FCALL_ showed a uniform cytosolic and nuclear distribution, the fluorescence ratio was ∼1. The minor Golgi distribution was signified by the Golgi: Nucleus fluorescence ratio of 1.3±0.1 (Fig. 5C-top box plot) before blue light (Fig. 5C-top box plot). It has been demonstrated that prenylation of CaaX sequence at the Ct of a protein alone is insufficient for proteins to interact with the plasma membrane (47–50). Therefore, it was unclear whether the observed primarily cytosolic localization of the protein is due to the masked *Phe-duo* in the Jα helical conformation of the prenylated protein or limited prenylation of the protein polypeptide, again due to the masked *Phe-duo* (39). When we exposed cells to blue light (445 nm) to activate the photoswitch while imaging Venus at 515 nm excitation and 542 ± 30 nm emission at 1 Hz frequency, a robust and Golgi-exclusive Venus recruitment was observed upon with a t_1/2_=2±1 s, which is also signified by the significantly increased Golgi: Nucleus fluorescence ratio (1.7±0.2) (one-way ANOVA: *F*_1,23_ = 44.393, *p* < 0.001) (Fig. 5C-Top, 5D-plot, and Movie S1). The Golgi recruitment of Venus accompanied a complementary reduction of cytosolic Venus fluorescence. The recruitment to the Golgi was confirmed using the co-localization of Venus with the trans-Golgi marker, GalT-dsRed (Fig. 5F and Movie S2). We further confirmed that this recruitment is Golgi targeted, by showing that the ER marker CFP-KDEL does not overlap with the Venus recruited regions upon blue light exposure (Fig. 5F). After the Venus fluorescence in Golgi reaches the steady state, termination of blue light resulted in dislodging of Golgi-bound iLID to near pre-blue light level (Fig. 5C-Top images, 5D, and Movie S1). Keeping the cells in the dark for ∼5 minutes allowed the complete reversal of Venus fluorescence to the cytosol (Movie S1). These observations suggested that a fraction of the prenylated protein remains cytosolic before blue light exposure. Unmasking the *Phe-duo* promotes its interaction with the Golgi membrane. Blue light termination associated *Phe-duo* masking disrupts this interaction. These observations validate our hypothesis that the prenyl anchor adjoining *Phe-duo* is crucial in promoting protein-membrane interactions. Next, to examine whether the masked *Phe-duo* in Venus-iLID_FCALL_ determines its prenylation potential during protein expression, we exposed cells to 450 nm blue LED light (5-second ON-OFF cycles for 12 hours in a CO_2_ incubator). The resultant cells showed a significant Golgi localization of Venus with Golgi: Nucleus of 1.6±0.1, which increased to 1.8±0.1, upon blue light exposure (one-way ANOVA: *F*_1,18_ = 4.893, *p* = 0.040) (Fig. 5C-middle). As a control experiment, we generated a Venus-iLID variant with an exposed *Phe-duo* by placing a linker sequence (GGGG) between iLID and *Phe-duo* (FF) (Venus-iLID_GGGGFFCALL_) (Fig. 5C-bottom). Compared to the cytosolic distribution observed upon expression of Venus-iLID_FCALL_, Venus-iLID_GGGGFFCALL,_ showed an enhanced Golgi and a detectable ER distribution (Fig. 5C-bottom). The Golgi: Nucleus Venus ratio (1.8±0.2) was the highest here, with very low Venus fluorescence in the nucleus. This is expected since in Venus-iLID_GGGGFFCALL_ expressing cells, prenyl-*Cys*-adjacent *Phe-duo* is exposed and free to interact with endomembrane. It was also not surprising that the blue light irradiation did not significantly change the distribution in Venus (Golgi: Nucleus 1.9±0.1) (one-way ANOVA: *F*_1,22_ = 1.759, *p* = 0.198) (Fig. 5C-bottom). These data indicated that, in addition to the prenyl group, membrane-accessible hydrophobic residues significantly enhance the membrane interaction of the engineered protein. Further, the decreasing presence of nuclear fluorescence from Fig. 5C top to bottom panels suggested that prenylated proteins do not enter the nucleus.

When cells are exposed to 10 µM GGTI286 during protein expression, Venus in both Venus-iLID_FCALL_ and Venus-iLID_GGGGFFCALL_ showed completely cytosolic distributions indicating these proteins are geranylgeranylation sensitive (Fig. 5E). Furthermore, the blue light irradiation did not recruit Venus to the Golgi in Venus-iLID_FCALL_ under GGTI286 treated conditions (Fig. 5E). To further confirm that geranylgeranylation is required for Golgi recruitment of this protein, we examined Venus-iLID_FCALL_ expressing cells under FTI, GGTI, or Fluvastatin-treated conditions. Both Fluvastatin-treated and GGTI-treated cells showed uniform Venus distribution throughout the cytosol and the nucleus, and failed to show blue light-induced Venus recruitment to Golgi (Fig. 5E top and S3-A). However, cells upon FTI treatment (only inhibits farnesylation) behaved similarly to control cells and exhibited blue light-induced Venus recruitment to Golgi (Fig. S3-A). To emphasize that blue light-induced Golgi recruitment of these two proteins depends on their prenylation state, we used C➔A mutant versions of iLID_FCALL_ and iLID_GGGGFFCALL_ (Venus-iLID_FAALL_ and Venus-iLID_GGGGFFAALL_) that showed complete cytosolic distribution before blue light exposure and no sensitivity to blue light irradiation (Fig. S3-B).

Overall, the data suggest that overnight blue light exposure during protein expression enhances the prenylation since the *Phe-duo* in the photoswitch is continuously exposed, improving polypeptide-membrane interactions. The significant nuclear localization of Venus in Venus-iLID_FCALL,_ cells (due to reduced prenylation) and the reduced presence of nuclear-Venus in Venus-iLID_GGGGFFCALL_ cells and overnight blue light exposed Venus-iLID_FCALL_ cells (due to enhanced prenylation) also collectively suggest that the unmasked *Phe-duo* enhances the prenylation efficacy. For instance, Venus-iLID_FCALL_ cells exposed to overnight blue light must have enhanced prenylation efficacy and thus a significantly higher concentration of Venus – 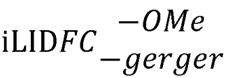 proteins compared to the control cells, as reflected by the significantly elevated Golgi localization of Venus. Collectively, these experiments demonstrate a previously unknown mechanism of polypeptide prenylation regulation. To our knowledge, this is the first presentation for dynamic control of molecular interactions using reversible optogenetics to regulate membrane interactions of lipidated proteins.

### 3.6 Contribution of pre-CaaX positively charged residues on Gγ prenylation efficacy and statin sensitivity is moderate, however significant

Both Gγ3-WT and Gγ9-WT contain homologous sequences consisting of positively charged *Lys, Arg* residues, or both at the beginning of their pre-CaaX regions (Fig. 3A). Our previous work indicated that the positively charged side chains of these residues help Gγ to maintain transient interactions with the phospholipid head groups of the membrane (41). To understand the collective role of hydrophobic and positively charged residues on enhanced membrane interactions of Gγ3-WT, we systematically mutated the pre-CaaX of Gγ9 to gradually achieve Gγ3-like characteristics without changing the prenylation type. Even though both *Lys* and *Arg* are positively charged, to examine whether Gγ3 achieved its improved membrane interactions and thereby resistance to Fluvastatin-induced inhibition of its membrane anchoring due to the higher geometric stability provided by *Arg* (51), we mutated the 61^st^ *Lys* in Gγ9 to *Arg* (Gγ9_KEK-GG <u>R</u>EK-GG_). It has been reported that compared to *Lys*, the guanidinium group of *Arg* allows the formation of stable and a larger number of electrostatic interactions in three different directions (51). The higher pKa of *Arg* may also contribute to forming more stable ionic interactions than *Lys* (51). Only ∼84% of the Gγ9_KEK-GG_→<u>_R_</u>_EK-GG_ mutant cells exhibited near-complete cytosolic distribution compared to ∼100% in Gγ9-WT (Fig. 6A, 6B and Table 3-Gγ9_KEK-GG→<u>R</u>EK-GG_). The appearance of a ∼16% cell population with partial cytosolic phenotype is also indicated by 45±9 s mobility t_1/2_, as opposed to the 9±5 s t_1/2_ of Gγ9 (Fig. 6C and Table 2-Gγ9_KEK-GG→<u>R</u>EK-GG_). These data showed that the pre-CaaX *Arg* grants Gγ3 a higher membrane affinity, thereby increasing prenylation efficacy and reducing statin sensitivity. Next, to understand the role of the additional *Lys* residue at the 69^th^ position of Gγ3-WT, we introduced an additional *Lys* to this mutant, creating the Gγ9_KEK-GG_→<u>_R_</u>_EK<u>K</u>GG_ mutant. Compared to both the wild type and Gγ9_KEK-GG_→<u>_R_</u>_EK-GG_ cells expressing Gγ9_KEK-GG_→<u>_R_</u>_EK<u>K</u>GG_ showed further reduced sensitivity to Fluvastatin (∼65% near-complete membrane anchorage inhibition) (Fig. 6A, 6B and Table 3-Gγ9_KEK-GG_→<u>_R_</u>_EK<u>K</u>GG_). This mutant also showed increased partial inhibition (∼36% cell) (Fig. 6B and Table 3-Gγ9_KEK-GG_→<u>_R_</u>_EK<u>K</u>GG_). These data indicate that basic pre-CaaX residues significantly enhance the prenylation efficacy while reducing the statin sensitivity of Gγ3. Building on this, we next generated a Gγ9 mutant (Gγ9_KEK-GG→KEK<u>KFF</u>_) to understand the cumulative effect of the additional *Lys69* and the *Phe-duo* in Gγ3 pre-CaaX. Compared to Gγ9-WT, Gγ9_KEK-GG→KEK<u>KFF</u>_ mutant cells exhibited elevated prenylation efficacy and reduced statin sensitivity, indicated by the significantly reduced near-complete membrane anchorage inhibition to ∼51%, and the increased partial inhibition to ∼48% (Fig. 6A, 6B and Table 3). This mutant only differs from the Gγ9_KEKGG→KEK<u>FF</u>_ mutant by having an additional *Lys,* which increases the partial inhibition from ∼16% to ∼51%. The sensitivity of these mutants to FTI, but not to GGTI, confirmed that their prenylation type is unchanged and remains farnesylated.

**Figure 6.**
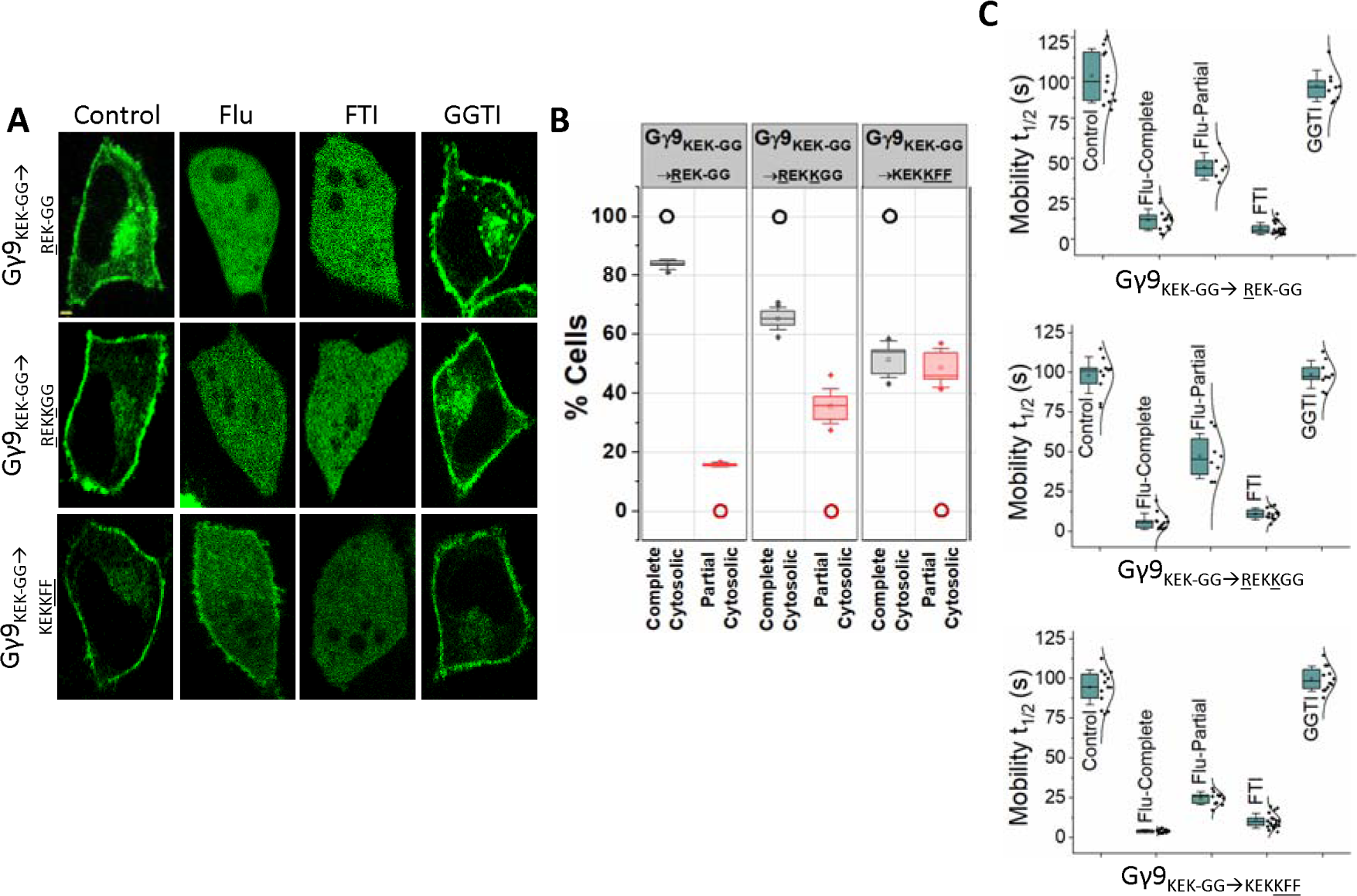
Positively charged pre-CaaX residues also contribute to Gγ prenylation efficacy and statin sensitivity. **(A)** Images of HeLa cells expressing GFP tagged Gγ9_KEK-GG_→<u>_R_</u>_EK-GG,_ Gγ9_KEK-GG_→<u>_R_</u>_EK<u>K</u>GG,_ and Gγ9_KEK-GG→KEK<u>KFF</u>_ mutants treated with vehicle (control), Fluvastatin (20 µM), FTI (1 µM), or GGTI (10 µM). Images represent the prominent phenotype observed in each population under given experimental conditions (scale: 5 μm; n ≥ 15 for each Gγ type). **(B)** Grouped box chart shows the percentages of cells in each Gγ mutant (Gγ9_KEK-GG_→<u>_R_</u>_EK-GG,_ Gγ9_KEK-GG_→<u>_R_</u>_EK<u>K</u>GG,_ and Gγ9_KEK-GG→KEK<u>KFF</u>_ mutants) showing near-complete (black) and partial (red) cytosolic distribution with Fluvastatin treatment. The black/red dotted circles indicate the % cells with near-complete or partial prenylation inhibition in wild-type Gγ9 expressing cells. **(C)** The whisker box plots show mobility half-time (t_1/2_) of proteins in (A) determined using fluorescence recovery after half-cell photobleaching. (Average box plots were plotted using mean±SD; Error bars: SD (Standard deviation); Gγ9_KEK-GG_→<u>_R_</u>_EK-GG_: n= 397 total number of cells from 5 independent experiments, Gγ9_KEK-GG_→<u>_R_</u>_EK<u>K</u>GG_: n= 764 total number of cells from 7 independent experiments, Gγ9_KEK-GG→KEK<u>KFF</u>_: n= 546 total number of cells from 5 independent experiments; Statistical comparisons were performed using One-way-ANOVA; p < 0.05; Flu: Fluvastatin; FTI: Farnesyl transferase inhibitor; GGTI: Geranylgeranyl transferase inhibitor).

## 4. Discussion

Post-translational modifications play central roles in membrane association of the G proteins such as Ras superfamily G proteins and heterotrimeric G proteins (33). Since G protein signaling pathways mediate a vast array of physiological responses, their dysregulation contributes to many diseases, including cancer, heart disease, hypertension, endocrine disorders, and blindness (52). In an early study, we showed that Gγ membrane interactions and associated signaling are inhibited in the presence of statins, likely due to the inhibition of prenylation. However, in the same study, we also observed differential extents of membrane interaction inhibition among different Gγ subtypes (13). In the present study, we attempted to identify the molecular reasoning for this differential inhibition.

Though prenylation is essential, it alone is insufficient for the plasma membrane interaction of prenylated G proteins (47–50). Studies show that in addition to the prenyl anchor, the amino acids antecedent to the CaaX motif further support membrane anchorage of proteins (40–42, 48, 49, 53, 54). However, in the absence of prenylation, the contributions of membrane-interacting Ct residues alone are not sufficient to support the recruitment of a protein to a membrane (13, 38). The positively charged poly *Lys* region on KRas-4b is a classic example showing the participation of prenyl-*Cys*-adjacent amino acids in targeting prenylated proteins to the plasma membranes (48, 49, 53). Here, KRas-4b _ membrane interaction is maintained by the thermodynamically favored insertion of the farnesyl lipid anchor into the membrane lipid bilayer, in concert with the electrostatic interactions of the 10 *Lys* residues (positively-charged) in the pre-CaaX region with the negatively charged polar headgroups of the plasma membrane phospholipids (53). It has also been shown that the polybasic residues in the pre-CaaX region of a protein can function as a sorting signal to prevent proteins from entering the Golgi (55). HRas is another example of containing a *Cys* residue adjacent to CaaX as a second signal to which a palmitoyl anchor is attached. The palmitoyl anchor on HRas serves as a second signal to traffic the protein to different targeted cell membranes (55). Along with the above evidence from previous literature, we have recently established that the *Phe-duo* adjacent to prenylated *Cys* in some Gγ subunits significantly promotes stronger membrane anchorage of Gγ (41). Two prenylation types and variable second signals encoded by the pre-CaaX region have been suggested to target a protein to different microdomains within the plasma membrane (55).

Even though the twelve Gγ types are prenylated with either one of the two prenyl anchors; 15-C farnesyl or 20-C geranylgeranyl, they exhibit a broad range of membrane affinities, spanning from the lowest membrane affinity observed in farnesylated Gγ9 to the highest in geranylgeranylated Gγ3 (40–42). When Gγs are prenylated, they reside at the inner leaflet of the plasma membrane as Gα_GDP_βγ heterotrimer, bound to or near GPCRs (56, 57). When GPCRs are activated, Gα exchanges the bound GDP to GTP, dissociating the heterotrimer into Gα_GTP_ and Gβγ. Upon GPCR activation, all Gγ members support Gβγ translocation from the plasma membrane to the endomembrane in a Gγ type-specific manner (40, 54). These Gγ type-dependent translocation rates and extents have been used as a measure of the membrane affinity of a specific Gγ (40–42, 54, 58–60). According to this classification, the three Gγs with the highest membrane affinities (Gγ2, γ3, and γ4) contain two conserved *Phe* residues that we named the *Phe-duo,* next to the prenylated and carboxymethylated *Cys* (41). An early study suggested that aromatic residues such as *Phe* in the membrane-interacting domain of a protein enhance membrane-protein interactions via a hydrophobic binding (61). This enhanced interaction is achieved by inserting the hydrophobic side chains into the fatty acid tail region of the lipid bilayer (61). Our recent work also indicated that the aromatic side chain of *Phe* residues is crucial for the enhanced membrane affinity of Gγ2, 3, and 4 (41). In contrast, the Gγ with the lowest membrane affinity, Gγ9, possesses two less hydrophobic *Gly* residues at the corresponding position. This lowest membrane affinity can easily be understood by considering Gγ9 prenylation type (farnesylation) and its less hydrophobic pre-CaaX region.

When the statin sensitivities of the 12 Gγ subtypes were examined by measuring the inhibition of membrane anchoring, we observed a partial perturbation of membrane anchorage in Gγ2, 3, 4, 7, 8, and 12. Interestingly, the same Gγ subtypes also exhibited a partial sensitivity to the GGTI. We propose the highly effective prenylation efficacy of Gγ2, 3, 4, 7, 8, and 12 allows them to achieve some prenylation by utilizing the residual geranylgeranyl transferase activity remaining under GGTI. These results also indicated that using the CaaX motif sequence to determine the type of prenylation is correct, and mobility t_1/2_ is a reliable measure of the G protein prenylation efficacy (62, 63). However, the complete to partial Fluvastatin sensitivity of geranylgeranylation-sensitive Gγs raised the question of whether this membrane anchorage inhibitions depends on the type of prenylation at all. To answer this question, we systematically mutated the CaaX and pre-CaaX regions of Gγ3 and Gγ9 since these two Gγs exhibited two extremities of Fluvastatin sensitivity out of 12 Gγ subtypes. The results from Gγ9_CALL_ and Gγ3_CIIS_ ruled out our hypothesis that Gγ sensitivity to statin is determined by the type of prenylation since both Gγ9_CALL_ and Gγ3_CIIS_ mutants retained their wildtype’s statin sensitivity despite the switched prenylation type. The significant change in statin sensitivity observed in Gγ9_Gγ3PC_ and Gγ3_Gγ9PC_ mutants further confirmed that Gγ’s sensitivity to statins is independent of prenylation type while highlighting a major role of pre-CaaX region in this process. Focusing on our early finding that *Phe-duo* in pre-CaaX regions adjacent to prenylated *Cys* plays a major role in membrane interactions of high membrane affinity Gγs, using Gγ9_GG→FF_ and Gγ3_FF→GG_ mutants, we next examined the influence of the *Phe-duo* on Gγ prenylation efficacy under statin-treated conditions. Our observations suggested that the hydrophobic residues in the pre-CaaX region control the prenylation efficacy of Gγ.

Our optogenetic approach that allowed reversible unmasking-masking of *Phe-duo* further emphasized the importance of *Phe-duo* on protein prenylation and membrane interaction of prenylated proteins. When the pre-CaaX FF is masked, we propose that the protein has a low prenylation efficacy. Indicating the multi-level regulation imposed, only a small fraction of the prenylated population binds Golgi before blue light, and pre-CaaX unmasking is required for its membrane binding. Although the exact reason is unclear, it has been shown that when other proteins do not support them, prenylated G proteins accumulate primarily at the Golgi, not in ER or the plasma membrane (64). Optical activation-induced Golgi recruitment of iLID model protein agrees with this. When the prenylated protein population is increased, we propose that the excess proteins Golgi cannot accommodate interact with the ER. This is evident from the major Golgi and minor ER distribution of the iLID model protein in cells exposed to blue light pulses overnight. In the iLID protein with GGGGFF pre-CaaX motif, the FF is constitutively unmasked, increasing prenylation. This protein shows both Golgi and ER localization.

Our observations upon altering positively charged residues of Gγ pre-CaaX, in which *Lys* and *Arg* influenced, however, differently, both the statin as well as the prenyl transferase inhibitor sensitivity is consistent with the expected contribution of basic residues on membrane interaction of a protein (65–67). Considering the positively charged pre-CaaX of Gγ9_Gγ3PC_ due to the presence of an *Arg* and an extra *Lys* in the pre-CaaX promoting membrane interactions, the similar G protein distributions observed for Fluvastatin-exposed Gγ9_Gγ3PC_ (despite its farnesylation) and Gγ3-WT can be understood. In summary, these data suggest that collectively both hydrophobic and positively charged residues in the pre-CaaX are crucial determinants of the statin sensitivity of a protein. Since prenylation takes place at the ER membrane, we also hypothesize that these pre-CaaX residues control the Gγ prenylation efficacy by contributing to Gγ polypeptide interactions with endomembrane and/or prenyltransferases during prenylation, post-prenylation processing, or both through their transient membrane interactions.

When the overall data summary is considered, it also appears that the contribution to the gain or loss of prenylation efficacy by a pre-CaaX residue is dependent on the type, number, location of other residues (composition of the pre-CaaX). When considering the relative roles of pre-CaaX and prenylation (and carboxymethylation), our data clearly shows that the latter is the major membrane recruiter, however, the efficacy of prenylation is pre-CaaX depedent, specially under suboptimal conditions. Regardless of their highly hydrophobic and/or positively charged pre-CaaX regions C➔A versions of these Gγ mutants exhibited completely cytosolic distributions and faster mobilities (Fig. S2), because pre-CaaX alone, without the prenyl anchor, cannot support membrane binding of a protein. To rule out bias in imaging data analysis, in addition to quantification of Gγ mobility with different treatment conditions, we carried out a blind control experiment with an experimenter who is blind to the type of mutation and the treatment condition. When compared, the results obtained by the blind control experimenter and the experimenter using one-way ANOVA, we did not observe a significant difference (Fig. S4). We further support our data by providing multi-cell images representing different phenotypes of each population of Gγ, WTs or their mutants in response to different treatments (Fig. S5 and 6).

In our previous study, we conclusively demonstrate that statins can differentially perturb Gβγ signaling in a Gγ type-specific manner by potentially inhibiting Gβγ-membrane interactions (13). The data from the current study provides a clear molecular explanation for this previously observed differential statin sensitivities, by revealing the collective role of hydrophobic and positively charged residues in the Gγ pre-CaaX region play. Given the cell and tissue-specific distribution of 12 Gγ types with distinct pre-CaaX motifs in the body, our data may provide a molecular justification for diverse pleiotropic effects of statins due to Gγ-dependent perturbation of Gβγ signaling (68–71). Additionally, our data also suggest that the extent of a G protein’s sensitivity to prenyltransferase inhibitors is also influenced by the nature of its pre-CaaX region. Due to the involvement of Ras family G proteins and some Gγ subtypes in oncogenesis (72–75), prenyltransferase inhibitors have been explored for chemotherapy (76). Tipifarnib, Lonafarnib, BMS-214662, L-778, and L-123 are a few farnesyltransferase inhibitors tested in phase-II clinical trials. Nevertheless, they failed, and the mutations acquired in the oncogene or cancer reaching the metastatic state were considered culprits (15, 77). We propose that examining the pre-CaaX composition of the oncogenic G proteins in specific tumors may allow revisiting these inhibitors for treating certain cancers.

## Conclusion

Switching the intact prenylation and statin sensitivity observed upon switching the CaaX motif between Gγ3 and Gγ9 indicated that the CaaX residues of the unprenylated polypeptide do not have a significant role governing protein-membrane interaction, as well as the prenylation efficacy. The perturbations we made in the pre-CaaX of Gγ indicate that residues provide varying, and specific contributions to either gain or loss of statin, as well as prenyl transferase inhibitor sensitivities depending on the properties, location, and the number on the pre-CaaX. For instance, the *Phe-duo* in Gγ9 pre-CaaX environment only has a lesser contribution from gaining the sensitivity compared to their role in Gγ3 pre-CaaX environment. Overall, the results indicate that despite the prenylation type, the molecular properties of the Gγ pre-CaaX region regulate their prenylation efficacy. When the prenylation process is suboptimal, either because of limited prenyl lipid substrate availability due to pharmacological (statins) or genetic constraints or limited prenyltransferase activity (due to prenyltransferase inhibitors), or both, our data clearly show that different Gγ types, depending on their pre-CaaX show distinct prenylation responses. This allows G protein-GPCR signaling in cells and tissues with particular Gγ types to be differentially regulated by statins and prenyltransferase inhibitors. Considering the involvement of G proteins in physiology and pathology, the discovered molecular interactions that govern the prenylation process may help design more efficient and cellular-genetic-makeup-dependent regulators for G protein signaling. All the prenyltransferase inhibitors have advanced only up to phase II clinical trials as cancer therapeutics. Additionally, our results also shed light on potential future combinatorial statin-transferase inhibitor therapies for a variety of diseases, including cancer.

## 2. Materials and methods

### 2.1 Reagents

The reagents used were as follows; Fluvastatin, Tipifarnib, and GGTI286 (Cayman Chemical, Ann Arbor, MI) were dissolved in appropriate solvents according to the manufacturer’s instructions and diluted in 1% Hanks’ Balanced Salt solution (HBSS) supplemented with NaHCO_3_ or regular cell culture medium before being added to cells.

### 2.2 DNA constructs and cell lines

For the engineering of DNA constructs used, GFP-Gγ9_Gγ3PC_, GFP-Gγ9_GG→FF_, Gγ3_FF→GG_, mCh-Gγ3_C72A_ YFP-tagged Gγ1–Gγ13, GFP-Gγ9, GFP-Gγ3, GFP-Gγ9_CALL_, GFP-Gγ3_CIIS_, CFP-KDEL, and GalT-dsRed have been described previously (13, 41, 54, 78). Dr. Brian Kuhlman from the University of North Carolina at Chapel Hill, North Carolina kindly provided permission to use iLID construct. Gγ3, Gγ9 mutants, and iLID constructs were generated by PCR amplifying the parent constructs in pcDNA3.1 (GFP-Gγ3, GFP-Gγ9, and Venus-iLID) with overhangs containing expected nucleotide mutations and DpnI (NEB) digestion (to remove the parent construct) followed by Gibson assembly (NEB) (79). Cloned cDNA constructs were confirmed by sequencing (Genewiz). The HeLa cell line was originally purchased from the American Type Culture Collection (ATCC) and authenticated using a commercial kit to amplify nine unique STR loci.

### 2.3 Cell culture and transfections

HeLa cells were cultured in minimum essential medium (MEM-Cellgro) with 10% heat-inactivated dialyzed fetal bovine serum (DFBS; from Atlanta Biologicals) and 1% penicillin−streptomycin (PS) in 60 mm tissue culture dishes and maintained in a 37°C, 5% CO_2_ incubator. When the cells reached ∼80% confluency, they were lifted from the dish using versene-EDTA (Cell-Gro) and resuspended in their growth medium at a cell density of 1 × 10^6^/ml. For imaging experiments (Gγ subcellular distribution analysis, fluorescence recovery after half-cell photobleaching for Gγ mobility analysis, subcellular distribution analysis, and blue light-induced membrane interaction analysis of Ct-modified iLID), cells were seeded on 35 mm cell culture–grade glass-bottomed dishes (Cellvis) at a density of 8 × 10^4^ cells. The day following cell seeding, cells were transfected with appropriate DNA combinations (YFP-tagged Gγ1–Gγ13: 0.8 µg, GFP tagged Gγ or Gγ mutants: 0.2 µg, Venus-iLID_FCALL_ and Venus-iLID_GGGGFFCALL_: 0.8 µg per each dish) using Lipofectamine 2000 transfection reagent (Invitrogen) according to the manufacturer’s protocol and stored in a 37°C, 5% CO_2_ incubator. After 3.5/5 hours of transfection, cells were replenished with the growth medium containing either dimethyl sulfoxide (DMSO) or the respective inhibitor (Fluvastatin, FTI, or GGTI). Live-cell imaging was performed ∼16 hours post-transfection.

### 2.4 Live cell imaging, half-cell photobleaching, iLID photoactivation, image analysis, and data processing

Live-cell imaging experiments were performed using a spinning disk (Dragonfly 505) XD confocal TIRF imaging system composed of a Nikon Ti-R/B inverted microscope with a 60X, 1.4 NA oil objective and iXon ULTRA 897BVback-illuminated deep-cooled EMCCD camera. Photobleaching and photoactivation of spatio-temporally confined regions of interest (ROIs) of cells were performed using a laser combiner with 40-100 mW solid-state lasers (445, 488, 515, and 594 nm) equipped with Andor® FRAP-PA unit (fluorescence recovery after photobleaching and photoactivation), controlled by Andor iQ 3.1 software (Andor Technologies, Belfast, United Kingdom). For selective photobleaching of YFP or GFP, 488 nm (3.1 mW) and 515 nm (1.5 mW) lasers at the focal plane (60x oil 1.49 NA objective) were used. Photobleaching speed: ∼100 µs per half cell (∼a 100 µm^2^ area). The photobleaching covers the entire depth of the cell along the Z axis. For photoactivation of iLID 445 nm laser at 6.3 µW was used. Proteins tagged with Venus/YFP were imaged using 515 nm excitation and 542 nm emission; GFP using 488 nm excitation−515 nm emission; CFP imaging or blue light activation of optogenetically active iLID constructs were performed using 445 nm excitation and 478 nm emission. For global and confined optical activation of iLID-expressing cells, the power of 445 nm solid-state laser was adjusted to 5 mW. Additional adjustments of laser power with 0.1%-1% transmittance was achieved using Integrated Lase Engine (ILE). Data acquisition, time-lapse image analysis, processing, and statistical analysis were performed as explained previously (13). Briefly, time-lapse images were analyzed using Andor iQ 3.1 software by acquiring the mean pixel fluorescence intensity changes of the entire cell or the selected ROIs.

### 2.5 Log cavity energy (Log CE) calculation Gγ Ct peptides (pre-CaaX+CaaX)

The hydrophobicity of a molecule can be quantified using octanol-water partition coefficient (*K_OW_*)-based Log Cavity energy (Log CE) value. The methodology to computationally determine Log CE using density functional theory (DFT) and the Solvent Model based on Density (SMD) was previously shown to have excellent agreement with experimental Log CE values for various molecules, including peptides (80, 81). We employed the M11 meta-functional (82) with SMD (83) and computed the Log CE using the equation, 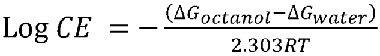 for the unprenylated Gγ polypetide (consisting pre-CaaX and CaaX regions) and prenylated and carboxymethylated Ct peptides representing the Gγ family members at 37 LJC (310 K). The peptides structures were optimized with a split-valance basis (3-21G*), and the solvation energies (Table S1) were computed with a polarized triple-zeta basis set (6-311+G**). All computations were performed using Gaussian16 (Revision C.01) software (84).

### 2.6 Computationally modeled structure generation of the iLID-based photoswitch

We used the amino acid sequence of iLID(46) to generate the modeled protein structures in Fig. 5A and B using AlphaFold2. First, we added the respective additional amino acids ‘FCALL’ and ‘GGGGFFCALL’ at the C terminus of the iLID to generate iLID_FCALL_ and iLID_GGGGFFCALL_. Then we fed these sequences to AlphaFold2 and built homology models. Structures that best fit the experimental iLID structure (PDB ID: 4WF0) were selected for protein folding validation. Finally, the protein preparation tool in Schrodinger Maestro13.3.121 was used to optimize the selected protein model. When generating the modeled 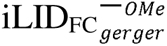 (before blue light irradiation), we used the 3D builder tool of Schrodinger and generated the carboxymethylated geranylgeranylated *cys* in 2D format. Then the structure was optimized using the protein preparation tool in Schrodinger Maestro for a low-energy 3D structure with corrected chirality. We followed a similar approach to generate the carboxymethylated geranylgeranylated Ct peptide 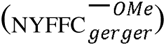 of light-activated 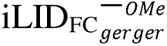 in 2D format, however used the Schrodinger Maestro LigPrep tool for optimizing the 3D structure. We then depicted AANDE residues of iLID as an unstructured peptide that connects AsLOV2 of iLID with the above prepared 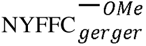. This structure is shown as the blue light activated protein in Fig. 5B.

### 2.7 Statistical data analysis

All experiments were repeated multiple times to test the reproducibility of the results. Statistical analysis and data plot generation were done using OriginPro software (OriginLab®). Results were analyzed from multiple cells and represented as mean ± SEM. The number of cells used in the analysis is given in respective figure legends. For Gαβγ heterotrimer mobility analysis using fluorescence recovery after half-cell photobleaching, after obtaining all the baseline-subtracted data employing the Nonlinear Curve Fitting (NLFit) tool in OriginPro, Gαβγ mobility dynamics plots were fitted to the ExpAssoc1 function under the Pharmacology category. For each fitting half time was calculated using, [half-time = *ln*(2)/K = *ln*(2)\**Tau*] equation where K is the rate constant and *Tau* is the time constant (*Tau* = 1/K). The mean values of half-time obtained from nonlinear curve fitting for all cells are given as mean mobility t_1/2_. One-way ANOVA statistical tests were performed using OriginPro to determine the statistical significance between two or more populations of signaling responses. Tukey’s mean comparison test was performed at the p < 0.05 significance level for the one-way ANOVA statistical test.

### 2.8 Blind control experiment

For each mutant with different treatment conditions, the blind control experimenter was given multiple images from experiments conducted on different days and instructed to categorize each cell image for its Gγ distribution, whether cytosolic or partially cytosolic. The mean values for each category of the blind control were compared with the values observed by the experimenter. The compared mean values were found not significantly different between the blind control experimenter and the experimenter (one-way ANOVA).

## Data availability

The datasets used and/or analyzed during the current study are available from the corresponding author upon reasonable request.

## Acknowledgment

We thank the Department of Biology, Saint Louis University, for providing instrumentation and other support. Ohio Supercomputer Center for providing access to high-performance computing clusters and Gaussian software for DFT calculations. We also thank the Saint Louis University Institute for Drug and Biotherapeutic Innovation for providing computational resources and access to Schrödinger software, with funding from the Saint Louis University Research Institute. We also thank Ajith lab members Sithurandi Ubeysinghe, Dhanushan Wijayaratna, Senuri Piyawardana, Chathuri Rajarathna, and Aditya Chandu for various experimental support, comments, and discussions.

## Author Contribution

M.T. generated most of the Gγ mutants and Venus-iLID_FCALL_ construct unless otherwise mentioned. W.T. generated GFP tagged Gγ9_KEK-GG_→<u>_R_</u>_EK-GG_, Gγ9_KEK-GG→<u>R</u>EK<u>K</u>GG_, Gγ9_KEK-GG→KEK<u>KFF</u>_, Gγ3_F70G_, Gγ3_F71G_ mutants and Venus-iLID_GGGGFFCALL_ construct. M.T. performed the majority of the experiments. W.T. performed the mobility t_1/2_ calculation experiment and data analysis for the Gγ3_C72A_ mutant and made Movie S1. W.T. computationally modeled structures in Fig 5A and B. M.T. analyzed the data and prepared the figures. J.L.P computationally calculated the log CE of pre-CaaX+CaaX peptides of unprenylated Gγ. M.T. and A.K. conceptualized the project and wrote the manuscript.

