## Supplemental Information for "CaaX-motif adjacent residues control G protein prenylation under suboptimal conditions"

**Supporting Information**

| **Table S1: Octanol-water partition coefficients (*K_OW_*)-based *Log Cavity Energy* (Log CE) calculation of Gγ pre-CaaX+CaaX polypeptides at 37 ^o^C (310 K)** | | | | |
| --- | --- | --- | --- | --- |
| **Gγ type** | **ΔG (octanol)** (kcal/mol) | **ΔG (water)** (kcal/mol) | **ΔG (o/w)** (kcal/mol) | **Log CE** |
| Gγ1 | 7.84 | 25.5 | -17.66 | 2.98 |
| Gγ2 | 2.14 | 26.75 | -24.61 | 4.15 |
| Gγ3 | 2.14 | 26.75 | -24.61 | 4.15 |
| Gγ4 | 2.58 | 28.66 | -26.08 | 4.39 |
| Gγ5 | 1.49 | 20.74 | -19.25 | 3.24 |
| Gγ7 | 4.15 | 24.63 | -20.48 | 3.45 |
| Gγ8 | 2.96 | 26.46 | -23.5 | 3.96 |
| Gγ9 | 7.66 | 22.23 | -14.57 | 2.45 |
| Gγ10 | 3.09 | 20.8 | -17.71 | 2.98 |
| Gγ11 | 8.1 | 23.49 | -15.39 | 2.59 |
| Gγ12 | 5.51 | 26.24 | -20.73 | 3.49 |
| Gγ13 | 6.8 | 24.39 | -17.59 | 2.96 |

**Figure S1**

Control	Flu	FTI	GGTI

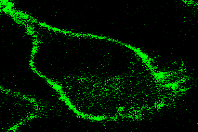

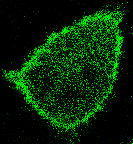

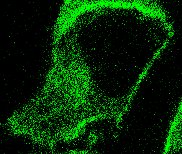

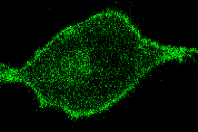

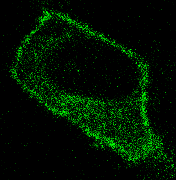

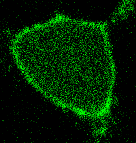

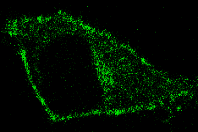

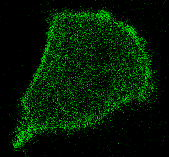

Gγ3_F70G_

Gγ3_F71G_

**A**

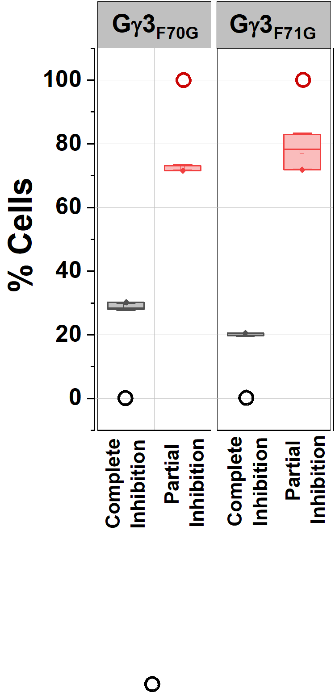

**B**

**C**

Gγ3_F70G_

Gγ3_F71G_

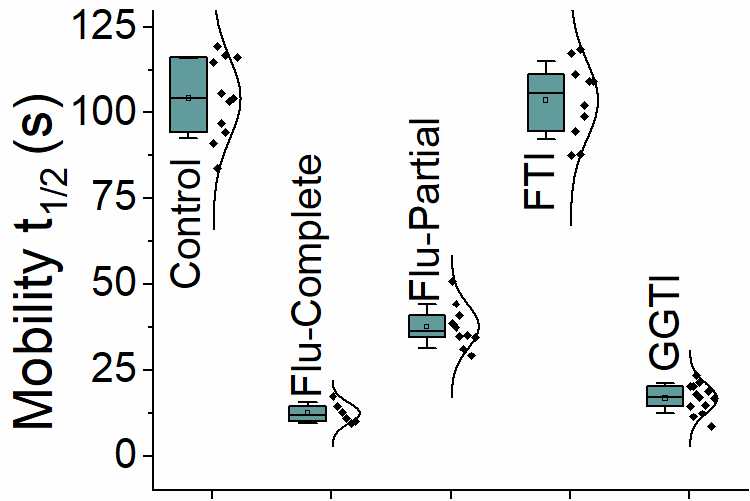

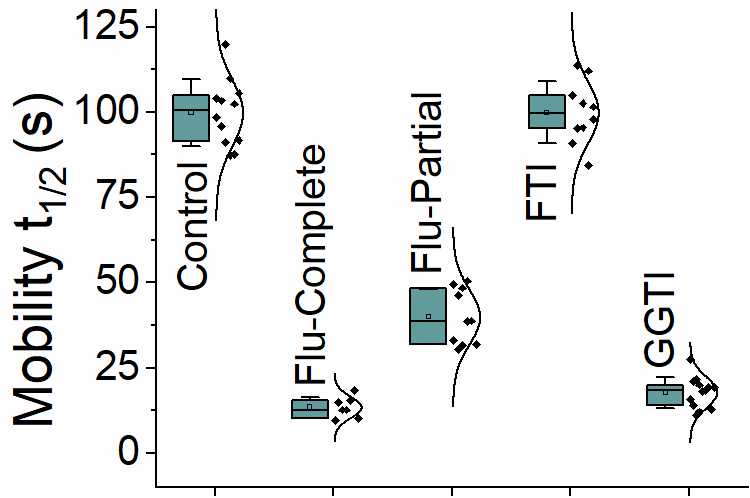

**Figure S1. Hydrophobicity of pre-CaaX residues determines Gγ prenylation efficacy and statin sensitivity.** **(A)** Images of HeLa cells expressing GFP tagged pre-CaaX mutants Gγ3_F70G_ and Gγ3_F71G_ exposed to the vehicle (control), Fluvastatin (20 µM), FTI (1 µM), or GGTI (10 µM). Images represent the prominent phenotype observed in each population under each experimental condition. Gγ9 mutants were sensitive to prenylation inhibition by FTI, while Gγ3 mutants were sensitive to GGTI-induced prenylation inhibition. Note the significance of having at least one *Phe* residue for prenylation in Gγ3 mutant compared to Gγ3_FF🡪GG_ mutant (Fig. 3B-bottom row) (scale: 5 μm; n ≥ 15 for each Gγ type). **(B)** Grouped box chart shows the percentages of cells in each Gγ mutant (Gγ3_F70G_ and Gγ3_F71G_ mutants) showing near-complete (black) and partial (red) cytosolic distribution with Fluvastatin treatment. The black/red circles indicate the % cells with near-complete or partial prenylation inhibition in each corresponding wild-type Gγ expressing cells (Gγ3_F70G_: n= 536 total number of cells, Gγ3_F71G_: n= 377 total number of cells; Each Gγ mutant was examined in 3 independent experiments) **(C)** The whisker box plots show the variations in mobility half time (t_1/2_) of G protein heterotrimers containing Gγ3_F70G_ and Gγ3_F71G_ mutants, treated with vehicle, Fluvastatin, FTI, or GGTI, determined using fluorescence recovery after half-cell photobleaching. (Average box plots were plotted using mean±SD; Error bars: SD (Standard deviation); Gγ3_F70G_: n= 49 total number of cells, Gγ3_F71G_: n= 50 total number of cells, all for mutants were examined in ≤ 3 independent experiments; Statistical comparisons were performed using One-way-ANOVA; p < 0.05; Flu: Fluvastatin; FTI: Farnesyl transferase inhibitor; GGTI: Geranylgeranyl transferase inhibitor).

**Figure S2**

Gγ3_CIIS_

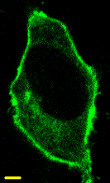

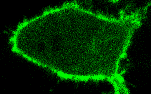

Gγ3_CIIS_

Flu

Gγ3_AIIS_

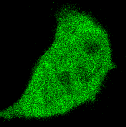

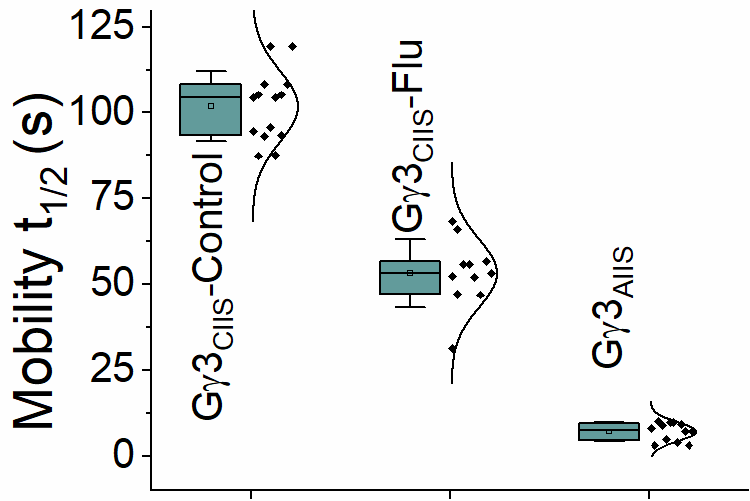

Gγ3_FF🡪LL_

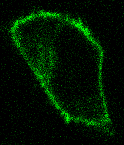

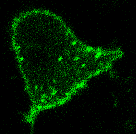

Gγ3_FF🡪LL_

Flu

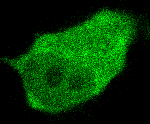

Gγ3_FF🡪LL(C🡪A)_

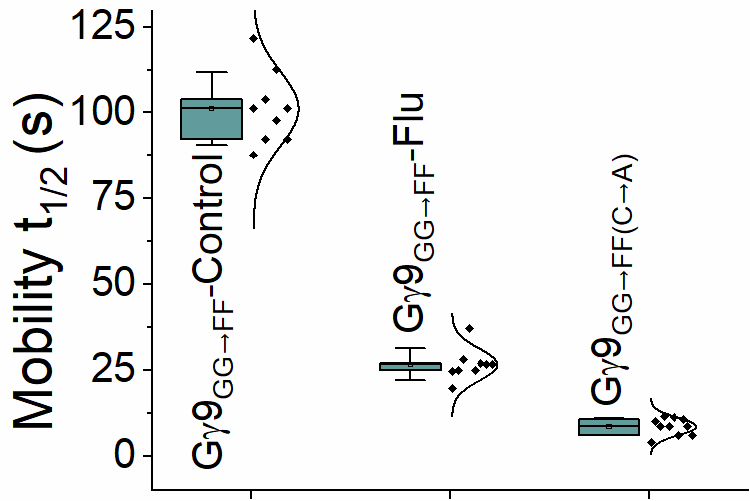

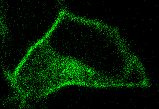

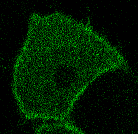

Gγ9_Gγ3PC_

Gγ9_Gγ3PC_-

Flu

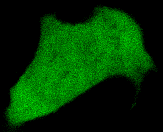

Gγ9_Gγ3PC(C🡪A)_

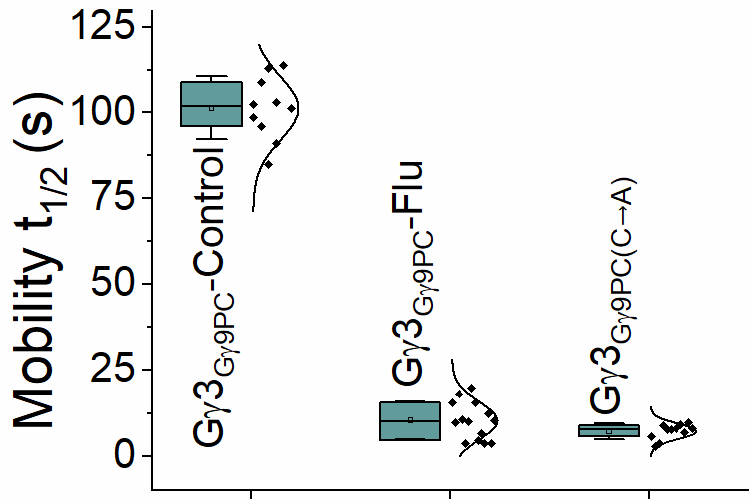

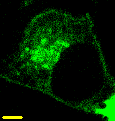

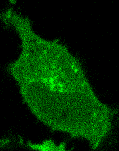

Gγ9_GG🡪FF_

Gγ9_GG🡪FF_

Flu

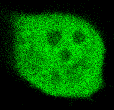

Gγ9_GG🡪FF (C🡪A)_

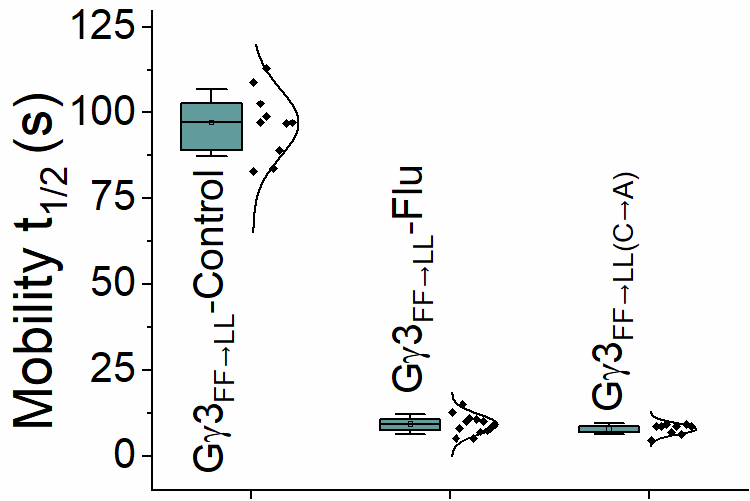

Gγ9_KEK-GG🡪_

_KEKKFF_

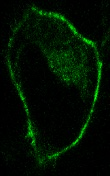

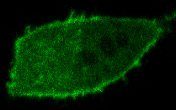

Gγ9_KEK-GG🡪_

_KEKKFF_

Flu

Gγ9_KEK-GG🡪_

_KEKKFF(C🡪A)_

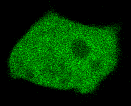

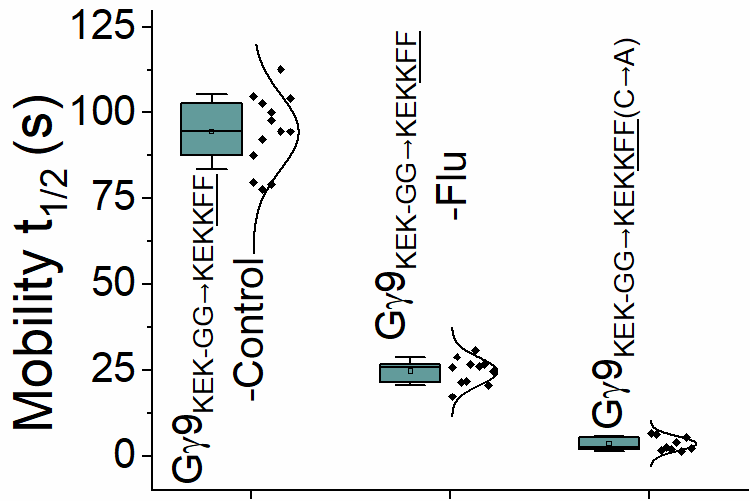

**Figure S2.** Images of HeLa cells expressing GFP statin treatment resisting Gγ mutants, their subcellular distribution upon Flu exposure (20 µM), and C🡪A mutated versions of each corresponding mutants. Images represent the prominent phenotype observed in each population under each experimental condition (scale: 5 μm; n ≥ 15 for each Gγ type). The whisker box plots show the variations in mobility half time (t_1/2_) of G protein heterotrimers containing each mutant determined using fluorescence recovery after half-cell photobleaching (Average box plots were plotted using mean±SD; Error bars: SD (Standard deviation); Statistical comparisons were performed using One-way-ANOVA; p < 0.05; Flu: Fluvastatin).

**Figure S3**

**(B)**

**(min)**

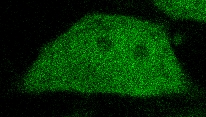

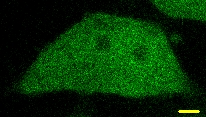

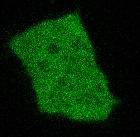

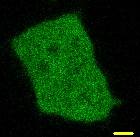

Venus iLID_FAALL_

Venus iLID_GGGGFFAALL_

Time	0	5

min

(min)

**No BL	BL**

**(A)**

**(min)**

Time	0	5

(min)

**No BL	BL**

**Venus-iLID_FCALL_**

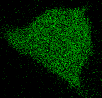

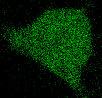

Flu

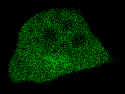

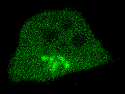

FTI

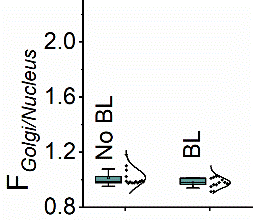

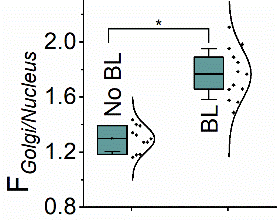

**Figure S3. (A)** Images of HeLa cells expressing Venus-iLID_FCALL_ showing an initial subcellular distribution of the protein upon Flu (20 µm) or FTI (1 µM) exposure and their change in subcellular distribution upon blue light irradiation (scale: 5 μm; n ≥ 15). The whisker box plots show the Golgi:Nucleus Venus fluorescence ratio under each condition (Average box plots were plotted using mean±SD; Error bars: SD (Standard deviation); Statistical comparisons were performed using One-way-ANOVA; p < 0.05; Flu: Fluvastatin FTI: Farnesyl transferase inhibitor ). **(B)** Images of Venus-iLID_FAALL_ and Venus-iLID_GGGGFFAALL_ showing subcellular distribution of the proteins before and after blue light irradiation (scale: 5 μm; n ≥ 15)_._

**Figure S4**

**Gγ3_FF🡪AA_**

**Gγ3_FF🡪YY_**

**Gγ3_FF🡪VV_**

**Gγ3_FF🡪GG_**

**Gγ9_GG🡪FF_**

**Gγ3_Gγ9PC_**

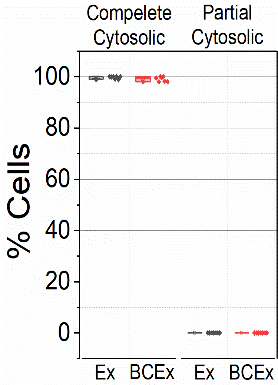

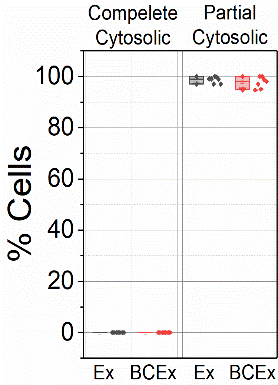

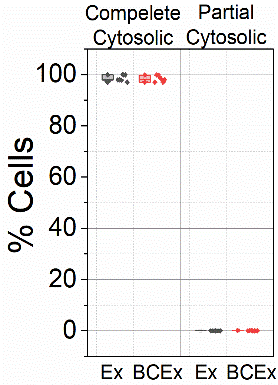

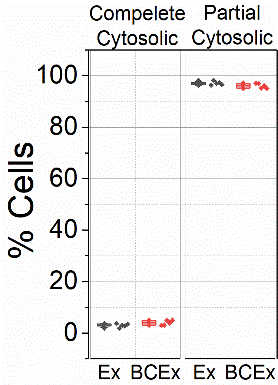

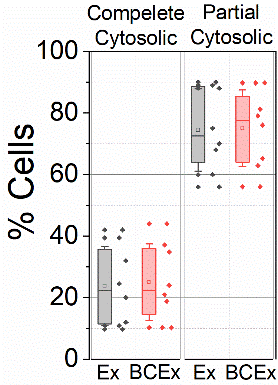

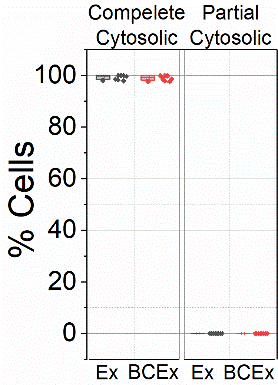

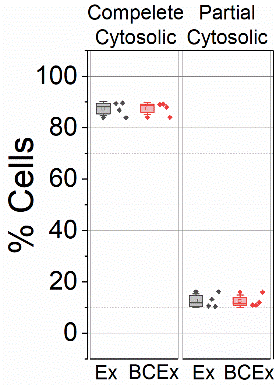

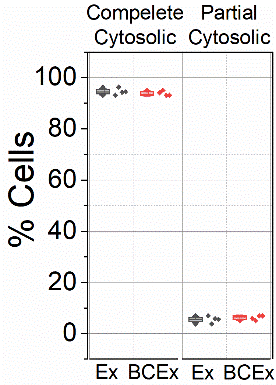

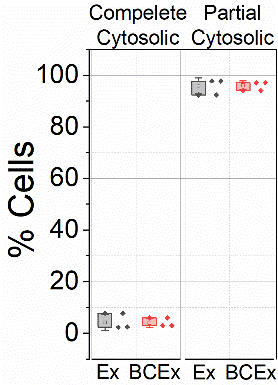

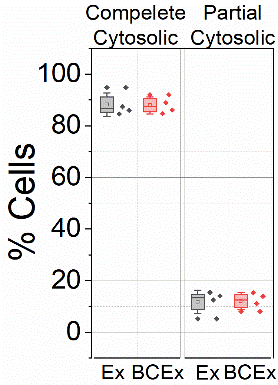

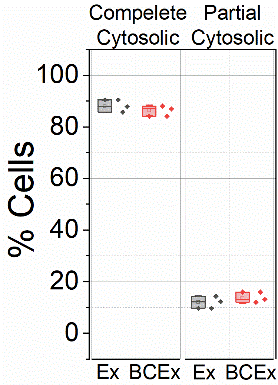

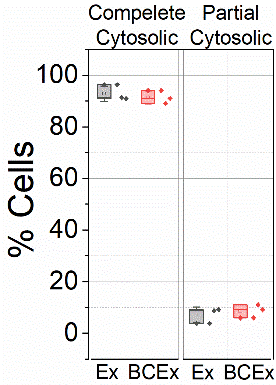

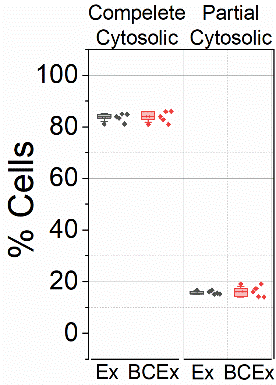

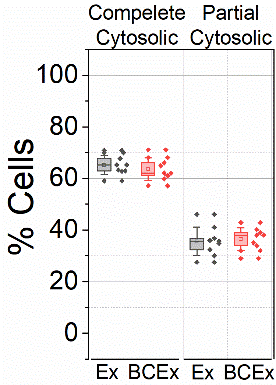

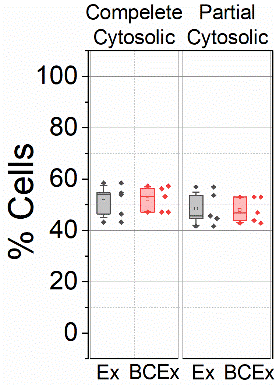

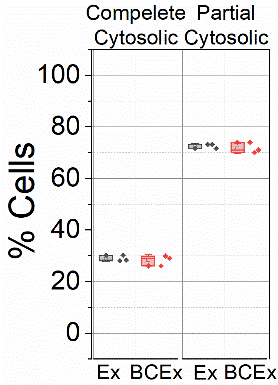

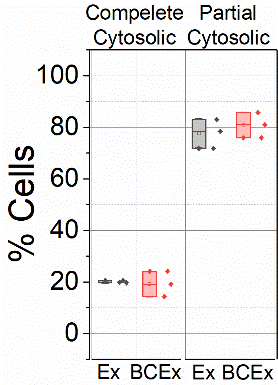

**Gγ9-WT**

**Gγ3-WT**

**Gγ9_CALL_**

**Gγ3_CIIS_**

**Gγ9_Gγ3PC_**

**Gγ3_FF🡪LL_**

**Gγ9_KEK-GG🡪 REK-GG_**

**Gγ9_KEK-GG🡪 REKKGG_**

**Gγ9_KEK-GG🡪 KEKKFF_**

**Gγ3_F70G_**

**Gγ3_F71G_**

**Figure S6: Summary of the blind control experiment results.** Grouped box charts show the percentages of cells in each Gγ mutant showing near-complete and partial cytosolic distribution with Fluvastatin treatment calculated by the experimenter (black) and the blind control experimenter (red) (Each Gγ mutant was examined in >3 independent experiments; mean±SD; Error bars: SD (Standard deviation); Statistical comparisons were performed using One-way-ANOVA; p < 0.05; Ex: Experimenter; BCEx: Blind Control Experimenter)

**Figure S5**

Gγ7 WT

Gγ11 WT

Gγ9 WT

Gγ10 WT

Gγ8 WT

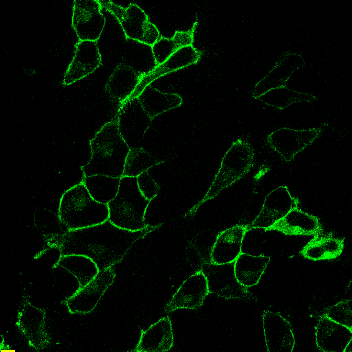

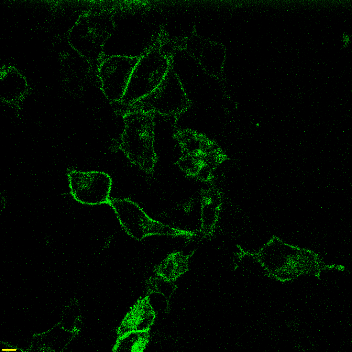

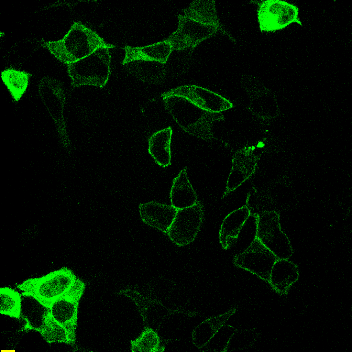

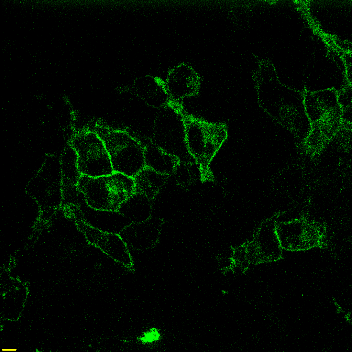

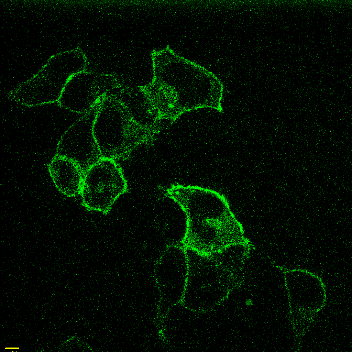

Control	Flu

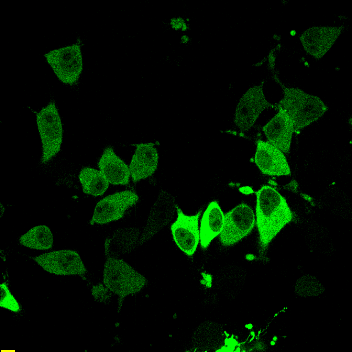

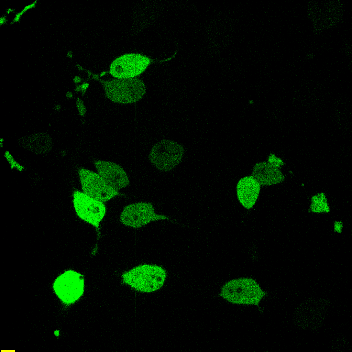

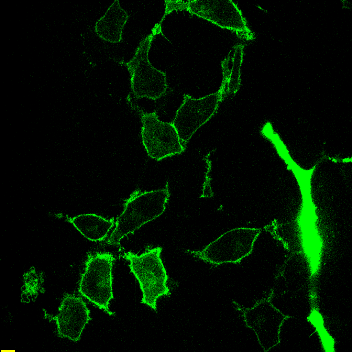

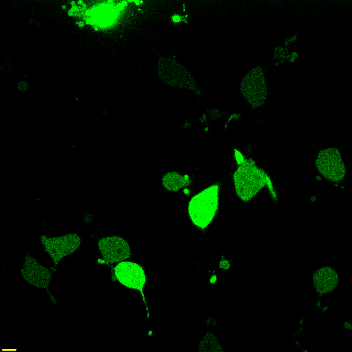

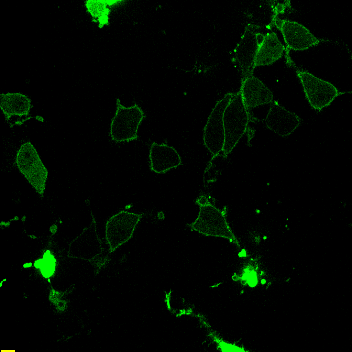

Control	Flu

Gγ1 WT

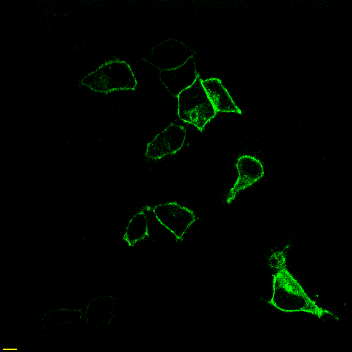

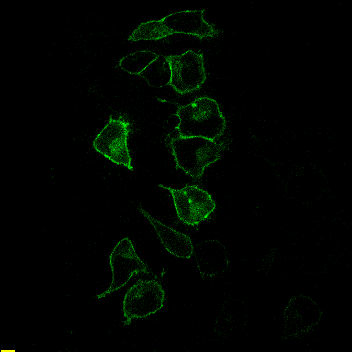

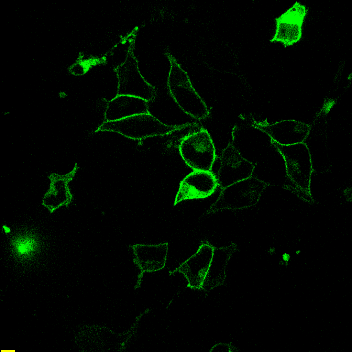

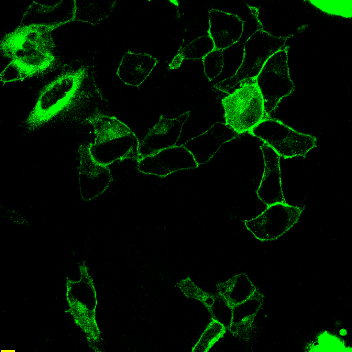

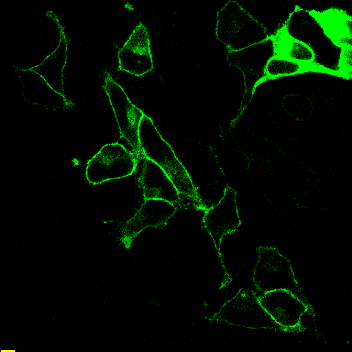

Gγ5 WT

Gγ3 WT

Gγ4 WT

Gγ2 WT

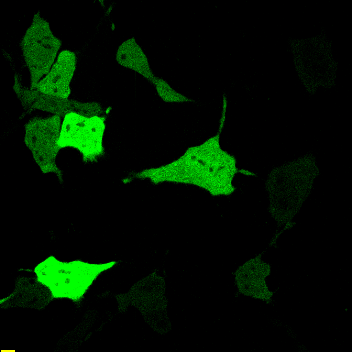

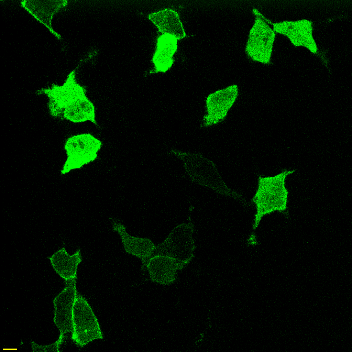

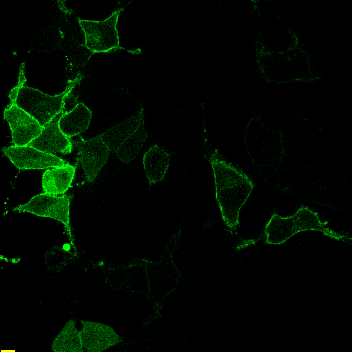

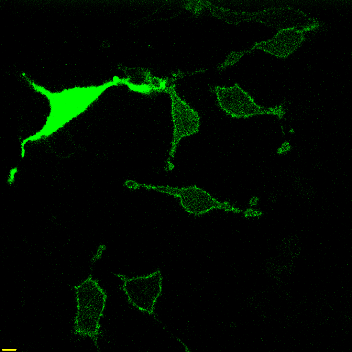

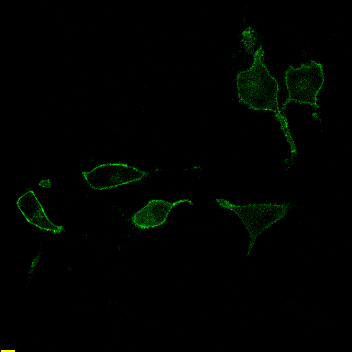

Control	Flu

Gγ12 WT

Gγ13 WT

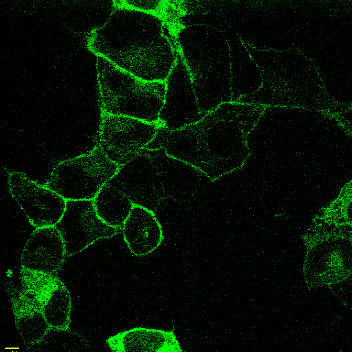

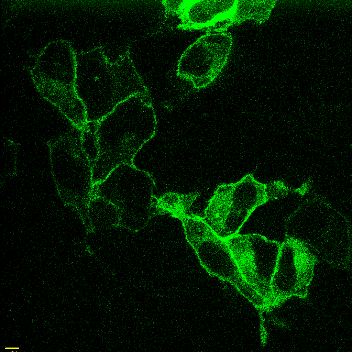

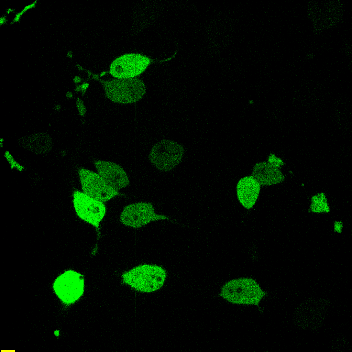

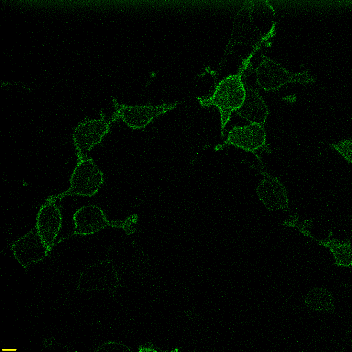

**Figure S5.** Multi-cell images representing different phenotypes of each population of Gγ under control and Flu (20 µm) treated conditions (scale: 10 μm)_._

**Figure S6**

Control	Flu

Gγ9_CALL_

Gγ3_CIIS_

Gγ9_Gγ3PC_

Gγ3_Gγ9PC_

Gγ9_GG🡪FF_

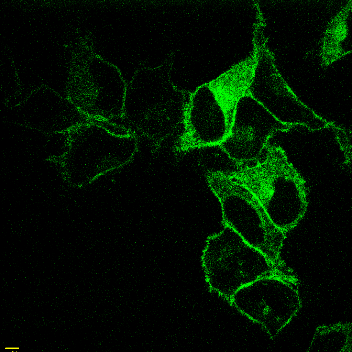

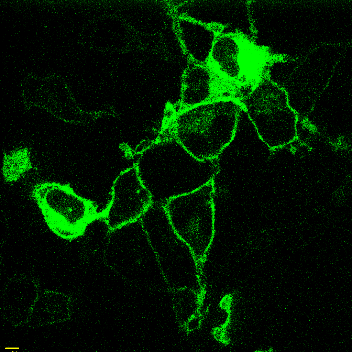

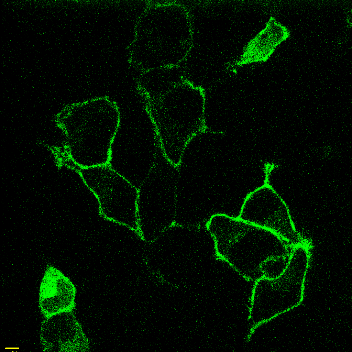

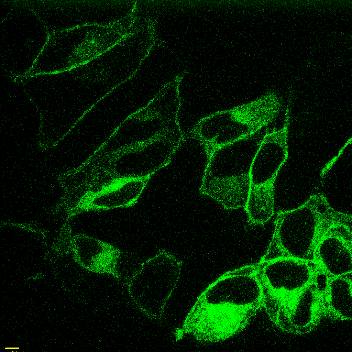

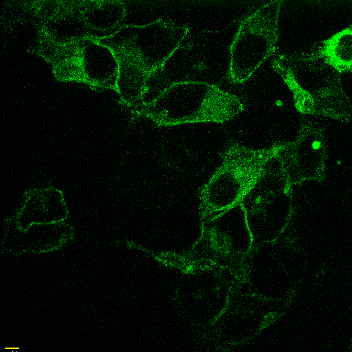

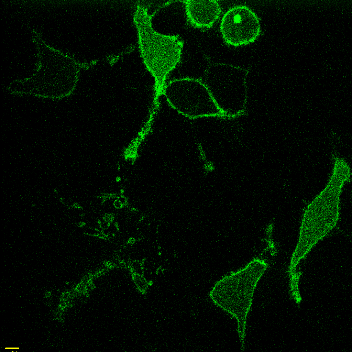

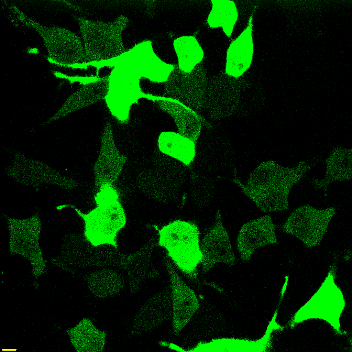

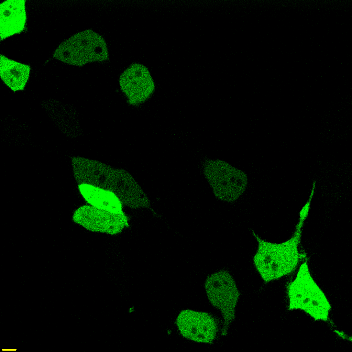

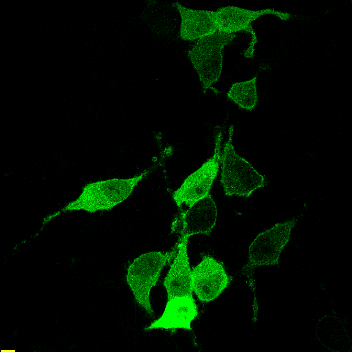

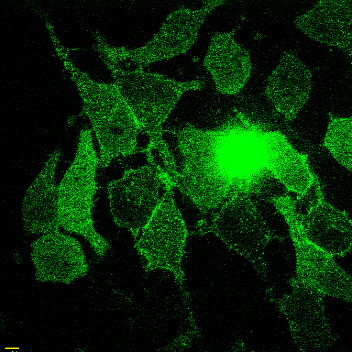

Gγ3_FF🡪LL_

Gγ3_FF🡪VV_

Gγ3_FF🡪YY_

Gγ3_FF🡪AA_

Gγ3_FF🡪GG_

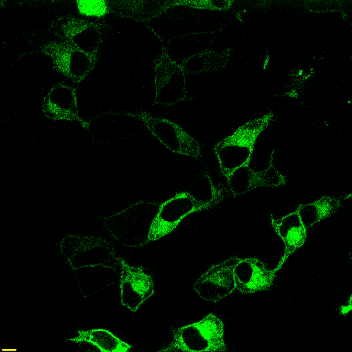

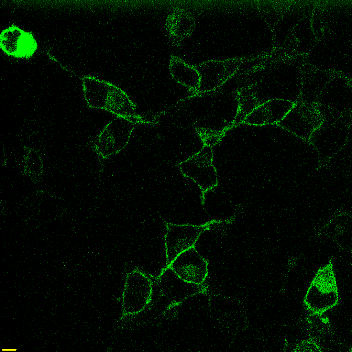

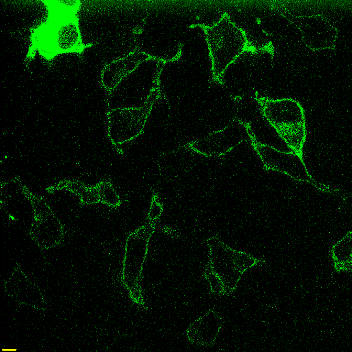

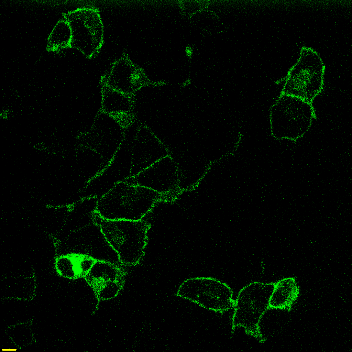

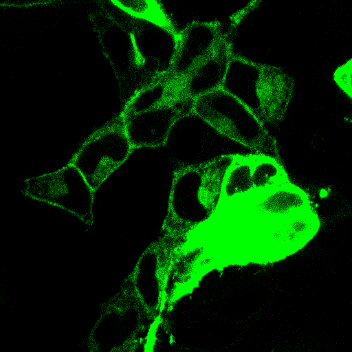

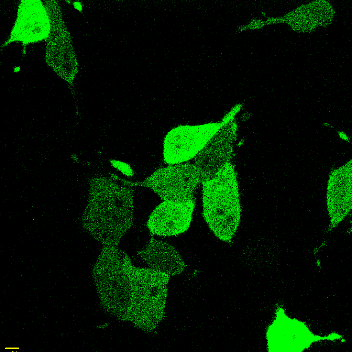

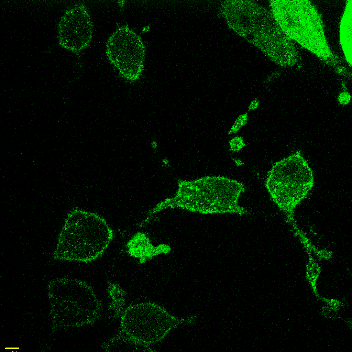

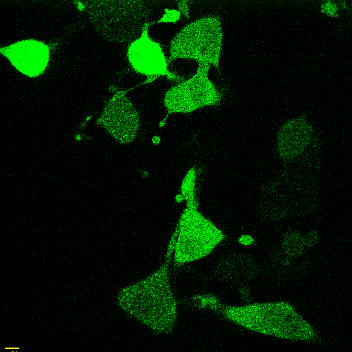

Control	Flu

Gγ9_KEK-GG🡪 REK-GG_

Gγ9_KEK-GG🡪_

_REKKGG_

Gγ9_KEK-GG🡪_

_KEKKFF_

Control	Flu

**Figure S6.** Multi-cell images representing different phenotypes of each population of Gγ mutant under control and Flu (20 µm) treated conditions (scale: 10 μm)_._

**Movie S1: Reversible optogenetic membrane affinity control of a prenylated protein.** A HeLa cell expressing Venus-iLID_FCALL_ shows the initial cytosolic distribution of the protein and its reversible recruitment to the Golgi upon blue light irradiation.

**Movie S2: Reversible optogenetic unmasking-masking of Ct hydrophobic residues governs reversible membrane recruitment of a prenylated protein.** 3-D images of HeLa cells expressing Venus-iLID_FCALL_ and GalT-dsRed show the initial cytosolic distribution of Venus fluorescence (pre-activation) and its Golgi recruitment with blue light irradiation (post-activation). Blue light-induced Golgi recruitment of Venus-iLID_FCALL_ is confirmed by its co-localization with the Golgi marker GalT-dsRed.
